# An epithelial *Nfkb2* pathway exacerbates intestinal inflammation by supplementing latent RelA dimers to the canonical NF-κB module

**DOI:** 10.1101/2020.11.02.365890

**Authors:** Meenakshi Chawla, Tapas Mukherjee, Alvina Deka, Budhaditya Chatterjee, Uday Aditya Sarkar, Amit K. Singh, Saurabh Kedia, Balaji Banoth, Subhra K Biswas, Vineet Ahuja, Soumen Basak

## Abstract

Aberrant inflammation associated with human ailments, including inflammatory bowel disease (IBD), is typically fuelled by the inordinate activity of RelA/NF-κB transcription factors. As such, the canonical NF-κB module mediates controlled nuclear activation of RelA dimers from the latent cytoplasmic complexes. What provokes pathological RelA activity in the colitogenic gut remains unclear. The noncanonical NF-κB pathway promotes immune organogenesis involving *Nfkb2* gene products. Because NF-κB pathways are intertwined, we asked if noncanonical signaling aggravated inflammatory RelA activity. Our investigation revealed frequent engagement of the noncanonical pathway in human IBD. In a mouse model, an *Nfkb2* function exacerbated gut inflammation by amplifying the epithelial RelA activity induced upon intestinal injury. Our mechanistic studies clarified that cell-autonomous *Nfkb2* signaling supplemented latent NF-κB dimers leading to hyperactive canonical RelA response in the inflamed colon. In sum, regulation of latent NF-κB dimers links noncanonical signaling to RelA-driven inflammatory pathologies and may provide for therapeutic targets.

**In brief:** Noncanonical NF-κB signals in intestinal epithelial cells supplement latent RelA dimers that, in turn, aggravated canonical NF-κB response in the colitogenic gut exacerbating intestinal inflammation.

**Highlights:**

- Human IBD involves the frequent engagement of the noncanonical NF-κB pathway.
- Mice deficient in the noncanonical signal transducer *Nfkb2* are resistant to experimental colitis.
- Noncanonical NF-κB signaling supplements latent RelA NF-κB dimers.
- Noncanonical NF-κB signaling amplifies canonical NF-κB response to TLR ligands.

## Introduction

Disruption of the intestinal barrier exposes tissue-resident cells, including intestinal epithelial cells (IECs), to luminal microbial contents setting off inflammation. Calibrated expressions of pro-inflammatory genes by tissue-resident cells limit local infections by orchestrating the recruitment and activation of effector immune cells, such as neutrophils, macrophages, and helper T cells, leading to the restoration of intestinal homeostasis. However, excessive inflammation provokes unabated activity of these effector cells, causing tissue damage in mice subjected to experimental colitis and contributing to the pathogenesis of human IBD, including ulcerative colitis (Friedrich et al., 2019; Kiesler et al., 2015; Kotas and Medzhitov, 2015).

Microbial substances signal through the canonical NF-κB module for activating the RelA:p50 transcription factor (Mitchell et al., 2016). In unstimulated cells, preexisting RelA dimers are sequestered in the latent cytoplasmic complexes by the inhibitory IκB proteins, the major isoform being IκBα. Canonical NF-κB signaling directs NEMO:IKK2 (alternately known as NEMO:IKKβ) mediated phosphorylation of IκBs, which are then degraded by the ubiquitin-proteasome system. Signal-induced degradation of IκBs liberates the bound RelA dimers into the nucleus, where they activate genes encoding pro-inflammatory cytokines and chemokines. Negative regulators of canonical signaling, including IκBα and A20, normally ensure a controlled RelA activity in physiological settings. In contrast, inordinate RelA response by tissue-resident cells culminates into non-resolving, pathological intestinal inflammation (Karrasch et al., 2007; Liu et al., 2017). Indeed, the severity of disease correlated with the extent of RelA activation in human IBD (Han et al., 2017). Whereas lessening canonical signaling using antisense oligo or peptide inhibitors mitigated experimental colitis in mice (Neurath et al., 1996; Shibata et al., 2007). What intensifies nuclear RelA activity in the colitogenic gut remains unclear.

The noncanonical NF-κB pathway mediates the nuclear accumulation of the RelB:p52 heterodimer (Sun, 2017). A select set of TNF receptor superfamily members, including lymphotoxin-β receptor (LTβR), induces noncanonical signaling, which stimulates NIK dependent phosphorylation of the NF-κB precursor p100, encoded by *Nfkb2*. Subsequent proteasomal processing of p100 generates the mature p52 subunit, which in association with RelB, translocates to the nucleus and induces the expression of immune organogenic genes.

Interestingly, genome-wide association studies identified *LTBR* and *NFKB2* as candidate genes linked to the susceptibility loci for human IBD (Liu et al., 2015). On the other hand, IEC-intrinsic deficiency of LTβR or NIK exacerbated chemically induced colitis in knockout mice (Macho-Fernandez et al., 2015; Ramakrishnan et al., 2019). It was suggested that LTβR protects against intestinal injury by driving the production of IL-23, which induces IL-22 mediated tissue repair. Dendritic cells (DCs) are the prime producers of IL-23 in the intestinal niche. Surprisingly, NIK’s depletion in DCs ameliorated colitis in mice (Jie et al., 2018). More so, biochemical studies indicated that genes encoding IL-23 subunits p19 and p40 are generic NF-κB targets and are not activated solely by RelB:p52 (Mise-Omata et al., 2007). Of note, LTβR and NIK also have functions beyond noncanonical NF-κB signaling (Boutaffala et al., 2015). It is less well understood if the noncanonical signal transducers encoded by *Nfkb2* per se contribute to the inflammatory pathologies observed in human IBD or in colitogenic mice. While global *Nfkb2^−/−^* mice were partially resistant to experimental colitis (Burkitt et al., 2015), the molecular mechanism and relevant cell types that link *Nfkb2* functions to gut inflammation remain obscure.

The canonical and noncanonical NF-κB pathways are intertwined at multiple levels (Shih et al., 2011). Proteasomal processing of p105, another NF-κB precursor encoded by *Nfkb1*, produces the mature p50 subunit that forms RelA:p50. While p105 processing is mostly constitutive, noncanonical signaling stimulates further p50 production (Yilmaz et al., 2014). Second, the noncanonical signaling gene *Nfkb2* represents a RelA target (Basak et al., 2008). Third, p100, in association with p105, forms high-molecular-weight complexes, which also sequester RelA in the cytoplasm (Savinova et al., 2009; Tao et al., 2014). Unlike canonical-signal-responsive, latent NF-κB complexes where preexisting RelA dimers are sequestered by IκBs, these high-molecular-weight complexes consist of monomeric RelA species bound to the individual NF-κB precursors. Finally, p52 interacts with RelA producing a minor RelA:p52 heterodimer (Banoth et al., 2015). Akin to RelA:p50, RelA:p52 is sequestered by IκBs, activated upon canonical signaling, and mediates pro-inflammatory gene expressions. This intertwining enables crosstalks between canonical and noncanonical NF-κB pathways. In addition to RelB regulation, noncanonical signaling also modulates canonical NF-κB response, both in the innate as well as the adaptive immune compartments (Almaden et al., 2014; Banoth et al., 2015; Chatterjee et al., 2016). We asked if the noncanonical pathway modulated RelA-driven inflammation in the colitogenic gut leveraging this interlinked NF-κB system.

Here, we present experimental evidence of frequent activation of the noncanonical *Nfkb2* pathway in IBD patients. In a mouse model, we ascertained that an IEC-intrinsic *Nfkb2* function exacerbated RelA-driven inflammation in the colitogenic gut. Our mechanistic analyses explained that tonic, tonic, noncanonical *Nfkb2* signaling supplemented latent RelA:p50 and RelA:p52 dimers by inducing simultaneous processing of both p105 and p100. The *Nfkb2*-dependent regulation of latent RelA dimers provoked a hyperactive canonical response in epithelial cells during the initiation of experimental colitis, aggravating intestinal inflammation. Finally, we established a tight association between heightened RelA activity and elevated processing of both the NF-κB precursors in human IBD. We argue that the regulation of latent NF-κB dimers by the noncanonical *Nfkb2* pathway provides for a therapeutic target in RelA-driven inflammatory pathologies.

## Results

### Engagement of the noncanonical NF-κB pathway in human IBD

To understand the molecular basis for the inordinate RelA activity associated with pathological intestinal inflammation, we biochemically analyzed colonic epithelial biopsies from thirty IBD patients suffering from ulcerative colitis (see STAR✪METHODS and Figure S1A-S1D). As controls, we utilized biopsies from ten, otherwise IBD-free, individuals ailing from hemorrhoids. We first measured the NF-κB DNA binding activity in nuclear extracts derived from these tissues by electrophoretic mobility shift assay (EMSA, see Figure 1A and Supplementary Figure S1A). Corroborating previous studies (Andresen et al., 2005), our investigation broadly revealed a heightened nuclear NF-κB (NF-κBn) activity in IBD patients as compared to the control cohort. Our shift-ablation assay suggested that this activity consisted mostly of RelA:p50 with a modest contribution of RelA:p52 (Figure 1B). Albeit at low levels, we also detected RelB complexes in some subjects, for example in patient #P29 (Figure 1B). Focusing on nuclear RelA (nRelA) activity in RelA-EMSA, we further ascertained that the heightened NF-κBn activity in IBD patients was mainly attributed by RelA (Figure 1C and Figure S1A). Quantitative immunoblot analyses of whole-cell extracts derived from colon biopsies revealed a reciprocal two-fold decrease in the median abundance of IκBα in IBD patients compared to controls indicating ongoing canonical signaling in the inflamed gut (Figure 1D and Figure S1B). Notably, IBD patients exhibited substantial variations in nRelA (Figure 1C). We catalogued IBD patients with nRelA level equal to or greater than one-and-a-half-fold of the median value in the control cohort as nRelA_high_; otherwise, they were designated as nRelA_low_. The median nRelA value for the nRelA_high_ subgroup comprising of twenty-three patients was 8.9; whereas the nRelA_low_ subgroup had a median of 4.6. Interestingly, IκBα levels were almost equivalently reduced in nRelA_high_ and nRelA_low_ patient subgroups (Figure 1E). These studies disclose disconnect between the intensity of canonical NF-κB signaling and the amplitude of nuclear RelA activity in IBD.

**Figure 1:**
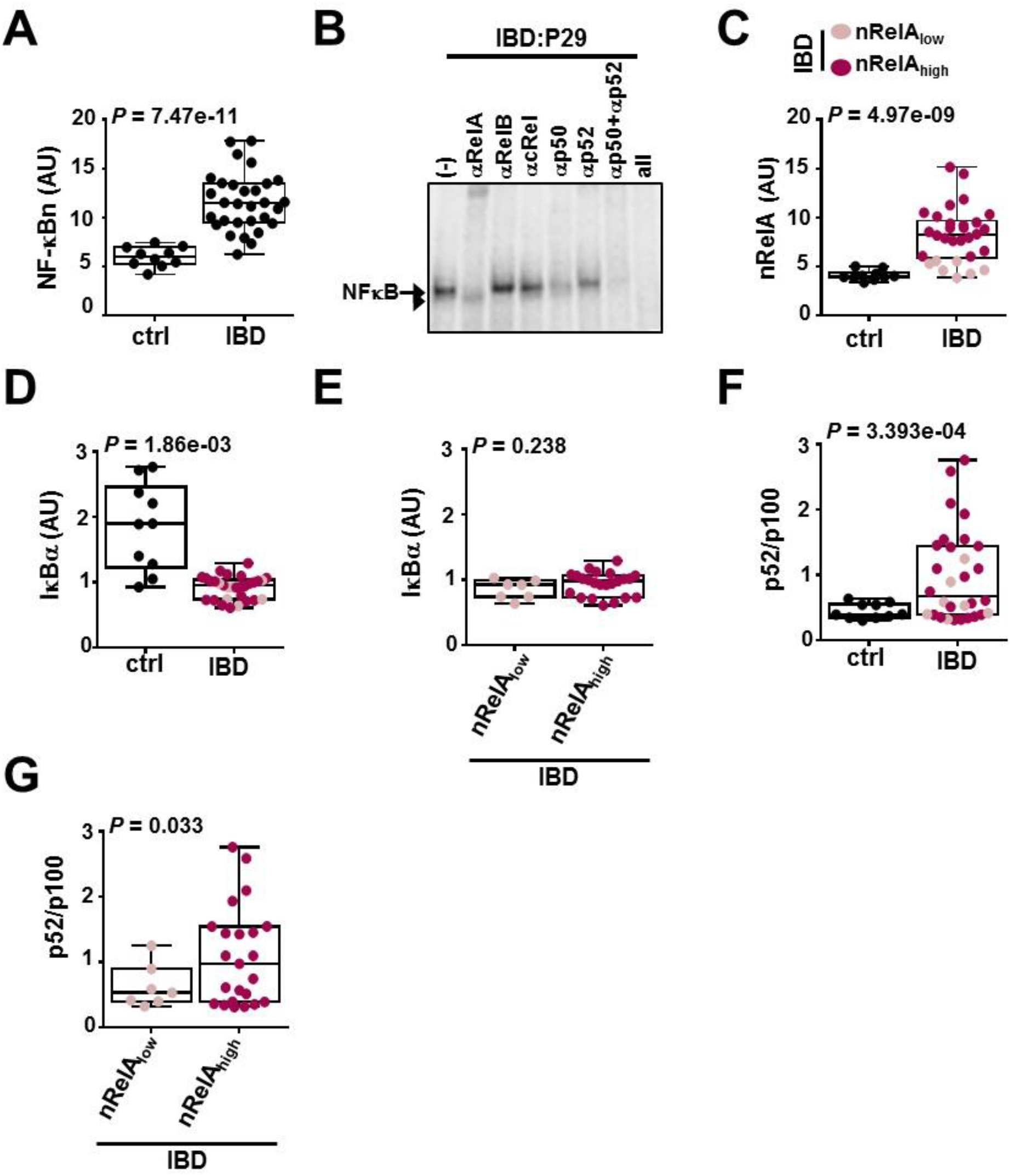
Heightened RelA activity correlates with elevated noncanonical NF-κB signaling in human IBD. **A.** and **C.** Nuclear extracts obtained using colon biopsies derived from ten non-IBD control patients and thirty IBD patients were examined by EMSA (see Figure S1). Signals corresponding to total (NF-κBn) (**A**) and RelA (nRelA) (**C**) NF-κB activities were quantified and graphed as a dot plot. Superimposed box plot indicates the median of the data along with first and third quartiles. **B.** The composition of nuclear NF-κB complexes present in colonic tissues was determined by shift-ablation assay. Antibodies against the indicated NF-κB subunits were used for ablating the respective DNA binding complexes in EMSA. A representative data has been shown. Arrow and arrowhead indicates RelA and RelB containing complexes, respectively. **D.** and **F.** Cell extracts obtained using colonic tissues from control individuals and IBD patients were examined by immunoblot analyses for the presence of IκBα and p52 as well as p100 (see Figure S1). Signals corresponding to IκBα in controls and IBD patients were quantified and graphed (**E**). Similarly, the abundance of p52 in relation to p100 was determined in these sets. **E.** and **G.** IBD patient subgroups with high or low nRelA activities in colonic tissues were compared for the abundance of IκBα (**E**) or the relative abundance of p52 to p100 (**G**). nRelA_low_ and nRelA_high_ patients have been color coded in other figure panels. In all panels, statistical significance was determined by Welch’s t-test on unpaired samples.

Next, we examined the plausible involvement of the noncanonical pathway in IBD. We measured the abundance of p52 in relation to p100 in colonic extracts as a surrogate of noncanonical signaling. Remarkably, up to 53% of IBD patients displayed elevated p100 processing, as determined from a one-and-a-half-fold or more increase in the p52:p100 ratio compared to the median p52:p100 value in the control cohort (Figure 1F and Figure S1B). Overall, IBD patients exhibited a more than two-fold increase in the median p52:p100 value. Interestingly, nRelA_high_ patients revealed significantly elevated p100 processing to p52 compared to nRelA_low_ patients (Figure 1G). Our cohort consisted of both steroid-naïve and steroid-treated individuals; however, steroid seemingly did not impact either the disease severity or NF-κB signaling at the time of biopsy acquisition (Figure S1C). We conclude that the noncanonical NF-κB pathway is frequently engaged in human IBD and that the strength of impinging noncanonical signaling correlates with the amplitude of nuclear RelA activity induced by the canonical NF-κB module in IBD patients.

### The noncanonical *Nfkb2* pathway amplifies epithelial RelA NF-κB response in the colitogenic murine gut

Because our investigation involving colonic epithelial biopsies implicated elevated noncanonical signaling in the heightened RelA activity observed in human IBD, we asked if the noncanonical *Nfkb2* pathway truly modulated epithelial RelA activation in the colitogenic gut. To address this, we treated mice with the colitogenic agent dextran sulfate sodium (DSS), collected IECs from these mice at various times post-onset of DSS treatment (Figure S2A), and subjected these cells to biochemical analyses. As such, DSS damages the mucosal layer triggering epithelial RelA activation via the canonical NF-κB module (Choi et al., 2015; Shibata et al., 2007). We found that WT mice elicited a strong NF-κBn activity at 12h post-onset of DSS treatment that persisted, albeit at a reduced level, even at 48h (Figure 2A). As observed with IBD biopsies, this activity consisted of mostly RelA:p50, and to some extent RelA:p52 (Figure 2B). Our immunoblot analyses further revealed a substantial basal processing of p100 into p52 in IECs derived from untreated WT mice; as suggested earlier (Allen et al., 2012), DSS treatment augmented this noncanonical *Nfkb2* signaling in the gut (Figure 2C). In response to DSS, *Nfkb2^−/−^* mice produced a rather weakened, but not completely abrogated, NF-κBn activity at 12h that further declined to a near-basal level by 48h (Figure 2A). Basal NF-κBn activity in untreated mice was comparable between these genotypes. We have previously shown that LTβR activates the noncanonical *Nfkb2* pathway in the gut (Banoth et al., 2015). Accordingly, intraperitoneal administration of an antagonistic LTβR-Ig antibody in mice diminished basal processing of p100 into p52 in IECs (Figure S2B). When subsequently challenged with DSS, mice administered with LTβR-Ig indeed produced a three-fold lessened NF-κBn activity indicating a role of tonic noncanonical signaling in shaping signal-induced canonical RelA response in IECs (Figure 2D).

**Figure 2:**
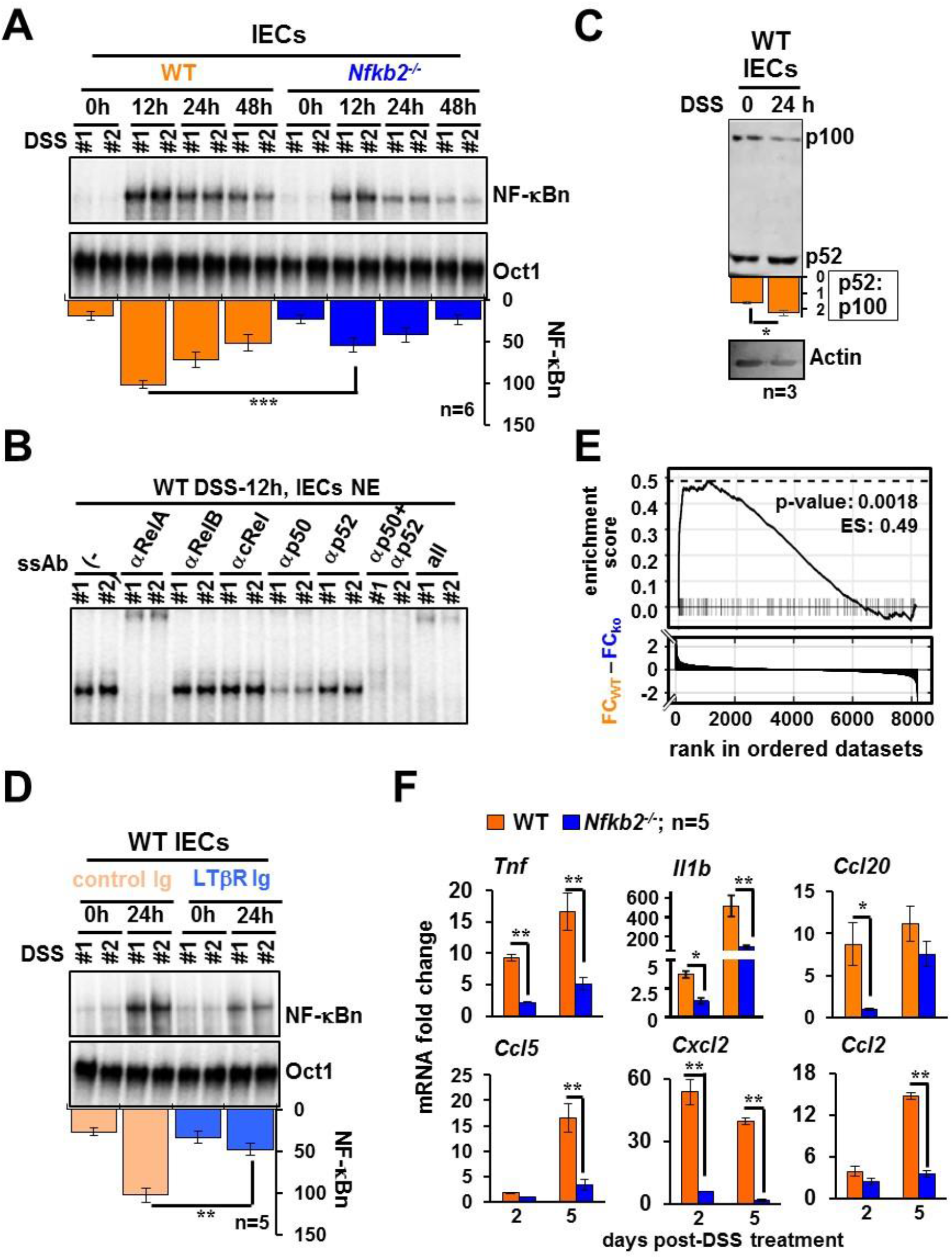
LTβR-*Nfkb2* signaling strengthens RelA NF-κB responses elicited by IECs in DSS-treated mice. **A.** WTand *Nfkb2^−/−^* mice were administered with 2.5% DSS. IECs were collected at the indicated times from the onset of DSS treatment and analyzed for NF-κBn activities. Bottom, NF-κBn signals were quantified and presented in the bargraphs below the respective lanes. **B.** Shift-ablation assay characterizing the composition of NF-κBn complexes induced in IECs of DSS-treated WT mice at 12h. Antibodies against the indicated NF-κB subunits were used for ablating the respective DNA binding complexes in EMSA. **C.** IECs collected from WT mice treated with 2.5% DSS were subjected to immunoblot analyses. Quantified signals corresponding to p52:p100 has been indicated. **D.** WT mice were intraperitoneally injected with control-Ig or LTβR-Ig, which blocks LTβR signaling, 24h prior to the onset of DSS treatment. Subsequently, IEC were isolated from mice treated with DSS for 24h and analyzed for NF-κBn by EMSA. **E.** GSEA comparing IECs derived from WT and *Nfkb2^−/−^* mice, either left untreated or treated with DSS for 48h (n=3 each), for RelA driven gene expressions. Briefly, RNA-seq analyses were performed and a list 8,199 genes with expressions in different data sets were prepared. The difference in the fold changes (FC) in the mRNA levels upon DSS treatment between WT and *Nfkb2^−/−^* mice was calculated for these genes, and genes were ranked in descending order of the fold change difference values (bottom panel). The relative enrichment of RelA target genes was determined by GSEA (top panel; ES denotes enrichment score. Each of the horizontal dashed line represents a RelA target gene. **F.** RT-qPCR revealing the abundance of indicated mRNAs in IECs derived from WT and *Nfkb2^−/−^* mice administered with DSS for two or five days. mRNA fold change values were calculated in relation to corresponding untreated mice. Quantified data represent means ± SEM. Two-tailed Student’s t-test was performed. \*\*\**P* < 0.001; \*\**P* < 0.01; \**P* < 0.05.

Pro-inflammatory gene activation by NF-κB factors in IECs represents a key event in the initiation of colitis (Friedrich et al., 2019; Mikuda et al., 2020). We performed RNA-seq analyses to address if the *Nfkb2* pathway tuned the transcriptional response of IECs in the colitogenic gut (STAR**✪**METHODS). A comparison of DSS-induced fold changes in the mRNA levels between WT and *Nfkb2^−/−^* mice revealed that *Nfkb2* deficiency led to both diminished inductions as well as hyperactivation of genes in IECs at 48h post-onset of treatment (Figure 2E and Figure S2C). Interestingly, gene set enrichment analysis (GSEA) of our transcriptomic data identified significant enrichment of RelA-targets among genes, whose DSS-responsive expressions in IECs were augmented by *Nfkb2* (Figure 2E). Our RT-qPCR analyses further demonstrated that the abundance of mRNAs encoding pro-inflammatory cytokines and chemokines was increased in IECs upon DSS treatment of WT mice (Figure 2F). While the levels of Ccl20 and Cxcl2 mRNAs were elevated within two days of the treatment, TNF, IL-1β, Ccl5, and Ccl2 mRNAs gradually accumulated over a period of five days. Importantly, *Nfkb2* deficiency restrained DSS-induced expressions of these RelA-target pro-inflammatory genes in IECs. The abundance of mRNA encoding TGFβ, which typically limits intestinal inflammation, was not considerably different between WT and *Nfkb2^−/−^* mice (Figure S2D). Our results indicate that by amplifying the epithelial RelA activity induced in the colitogenic gut, the tonic noncanonical *Nfkb2* pathway aggravates RelA-driven inflammatory gene program in mouse.

### A stromal *Nfkb2* function exacerbates experimental colitis in mice

In a recent study, Lyons et al. (2018) quantitatively assessed the contribution of IECs in experimental colitis (Lyons et al., 2018). Their investigation suggested that pro-inflammatory gene expressions by particularly undifferentiated IECs play a pivotal role in the infiltration of inflammatory immune cells in the colitogenic gut aggravating tissue injuries. Following those lines, we enquired if limiting RelA-driven gene response in IECs altered the course of DSS-induced ulcerating colitis in *Nfkb2^−/−^* mice. First, we collected cells from the colonic lamina propria of mice subjected to DSS treatment at day five and determined the composition effector immune cell subsets by flow cytometry (Figure S3). In comparison to WT mice, *Nfkb2^−/−^* mice exhibited less marked accumulation of inflammatory cells in the gut upon DSS treatment (Figure 3A). DSS-treated *Nfkb2^−/−^* mice displayed a five-fold reduced frequency of macrophages - identified as CD11b+F4/80+ cells, a close to two-fold decrease in the abundance of DCs - broadly considered as CD11c+F4/80-cells, and a two-and-a-half-fold reduced frequency of CD4+ T cells. Further investigation revealed a substantially lower colonic abundance of IFNγ+ Th1 cells and IL17A+ Th17 cells in DSS-treated *Nfkb2^−/−^* mice. However, DSS caused a more profound accumulation of suppressive CD4+CD25+FoxP3+ Treg cells in *Nfkb2^−/−^* mice presumably owing to their increased stability in the context of timid gut inflammation. The frequency of neutrophils - identified as Gr1+SiglecF-cells - was not discernibly different between WT and *Nfkb2^−/−^* mice subjected to DSS treatment. Untreated WT and *Nfkb2^−/−^* mice presented almost equivalent frequencies of these immune cells in the gut (Figure 3A).

**Figure 3:**
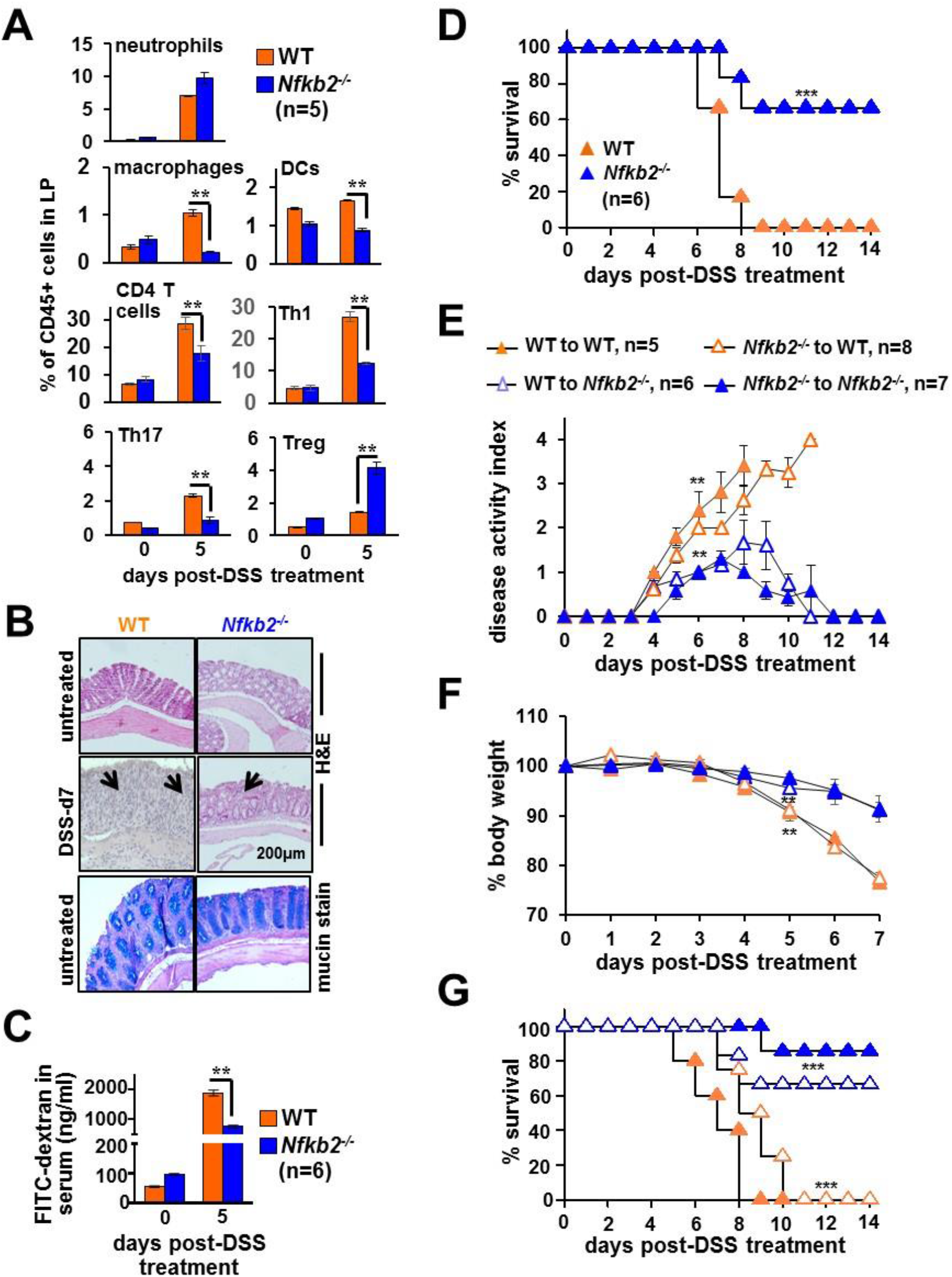
*Nfkb2* deficiency in the stromal compartment ameliorates chemically induced colitis in mice. **A.** Bar plot revealing relative frequencies of the indicated immune cells among CD45.2+ cells present in the lamina propria (LP) of WT and *Nfkb2^−/−^* mice. Mice were either left untreated or administered with 2.5% DSS. The composition of LP cells was examined by flow cytometry. **B.** Representative image showing H&E stained colon sections derived from untreated or DSS-treated mice of the indicated genotypes (top two panels). Colon sections from untreated mice were additionally stained using Alcian Blue (bottom panel). The data represent n=4; four fields per section and a total of five sections from each set were examined. Panels show 20X magnification. **C.** Bar chart revealing the concentration of FITC-dextran in serum of untreated or DSS-treated WT and *Nfkb2^−/−^* mice. FITC-dextran was gavaged 6h prior to serum collection. **D.** WT and *Nfkb2^−/−^* mice were administered with 2.5% DSS for seven days and monitored for survival for fourteen days. **E.** to **G.** Reciprocal bone marrow chimera generated using WT and *Nfkb2^−/−^* mice were subjected to 2.5% DSS treatment and then evaluated for the disease activity **(E)**, bodyweight changes **(F)**, and mortality **(G)**. Statistical significance was determined by comparing WT to *Nfkb2^−/−^* chimeras with WT to WT chimeras or *Nfkb2^−/−^* to WT chimeras with *Nfkb2^−/−^* to *Nfkb2^−/−^* mice. Quantified data represent means ± SEM. For Figure 3D and Figure 3G, the statistical significance was determined using log-rank (Mantel–Cox) test. Otherwies, two-tailed Student’s t-test was performed. \*\*\**P* < 0.001; \*\**P* < 0.01; \**P* < 0.05.

We then compared WT and *Nfkb2^−/−^* mice for DSS-inflicted pathologies. Our histological analyses revealed that colons from untreated WT and *Nfkb2^−/−^* mice were largely indistinguishable with respect to epithelial architecture and mucin expressions (Figure 3B). More so, DSS treatment for thirty-six hours caused almost equivalent IEC apoptosis indicating early epithelial injury of the similar extent in these genotypes (Figure S4A). At day seven, however, WT mice displayed extensive disruption of the epithelial barrier and widespread infiltration of leukocytes in the submucosa, while *Nfkb2^−/−^* mice exhibited less pervasive intestinal damage (Figure 3B). We also evaluated the intestinal barrier permeability by scoring serum concentrations of FITC-dextran gavaged orally to DSS-treated mice. Consistent with histological studies, DSS treatment caused only a modest increase in the intestinal permeability in *Nfkb2^−/−^* mice at day five (Figure 3C). WT mice subjected to acute DSS treatment also exhibited profound disease activity in these later days that was accompanied by a significant shortening of the colon and a close to fifteen percent loss in the bodyweight (Figure S4B-S4D). As reported (Burkitt et al., 2015), these colitis phenotypes were less pronounced in *Nfkb2^−/−^* mice (Figure S4B-S4D). Indeed, the entire WT cohort succumbed to DSS-induced colitis by day eight post-onset of DSS treatment, while two-thirds of *Nfkb2^−/−^* mice survived the course of colitis (Figure 3D).

*Nfkb2* signaling in the hematopoietic compartment curtails the generation of Treg cells (Dhar et al., 2019; Grinberg-Bleyer et al., 2018), while our global *Nfkb2^−/−^* mice displayed an increased frequency of Tregs in the colitogenic gut. Therefore, we further examined reciprocal bone marrow chimeras generated using WT and *Nfkb2^−/−^* mice for dissecting the hematopoietic and the stromal *Nfkb2* functions in experimental colitis. Regardless of WT or *Nfkb2^−/−^* hematopoietic cells, WT recipients were sensitive to DSS-induced colitis, exhibiting heightened disease activity, substantial bodyweight loss, and mortality (Figure 3E-3G). On the other hand, *Nfkb2^−/−^* mice showed resilience, even in the presence of WT hematopoietic cells. We conclude that a stromal *Nfkb2* function exacerbates experimental colitis by directing inflammatory infiltrates to the gut and that *Nfkb2* signaling in the hematopoietic compartment is less consequential for DSS-inflicted intestinal pathologies.

### An IEC-intrinsic role of *Nfkb2* aggravates experimental colitis in mice

Next, we asked if *Nfkb2* signaling in IECs was sufficient for aggravating intestinal inflammation. To test this, we generated *Nfkb2^ΔIEC^* mice, which specifically lacked *Nfkb2* expressions in IECs (Figure S4E). As compared to control *Nfkb2^fl/fl^* mice possessing otherwise functional floxed *Nfkb2* alleles, *Nfkb2^ΔIEC^* mice elicited a relatively moderate NF-κBn activity in IECs upon DSS treatment (Figure 4A). We consistently noticed a subdued expression of RelA-target pro-inflammatory genes in IECs of *Nfkb2^ΔIEC^* mice at day five of the DSS regime (Figure 4B). Our flow cytometry analyses further revealed a reduced accumulation of inflammatory immune cells, including macrophages, Th1, and Th17 cells, in the colonic lamina propria of DSS-treated *Nfkb2^ΔIEC^* mice (Figure 4C). Finally, a dysfunctional *Nfkb2* pathway in IECs alleviated experimental colitis – DSS treatment led to less severe disease activity and only a marginal loss of the bodyweight in *Nfkb2^ΔIEC^* mice (Figure 4D and Figure 4E). In contrast to DSS-inflicted mortality in *Nfkb2^fl/fl^* mice, most of the *Nfkb2^ΔIEC^* mice survived acute DSS treatment (Figure 4F). Our results substantiate that an IEC-intrinsic *Nfkb2* pathway exacerbates experimental colitis by amplifying RelA-driven inflammatory gene responses in IECs. In other words, we establish the functional significance of *Nfkb2*-mediated modulation of epithelial canonical signaling in aberrant intestinal inflammation.

**Figure 4:**
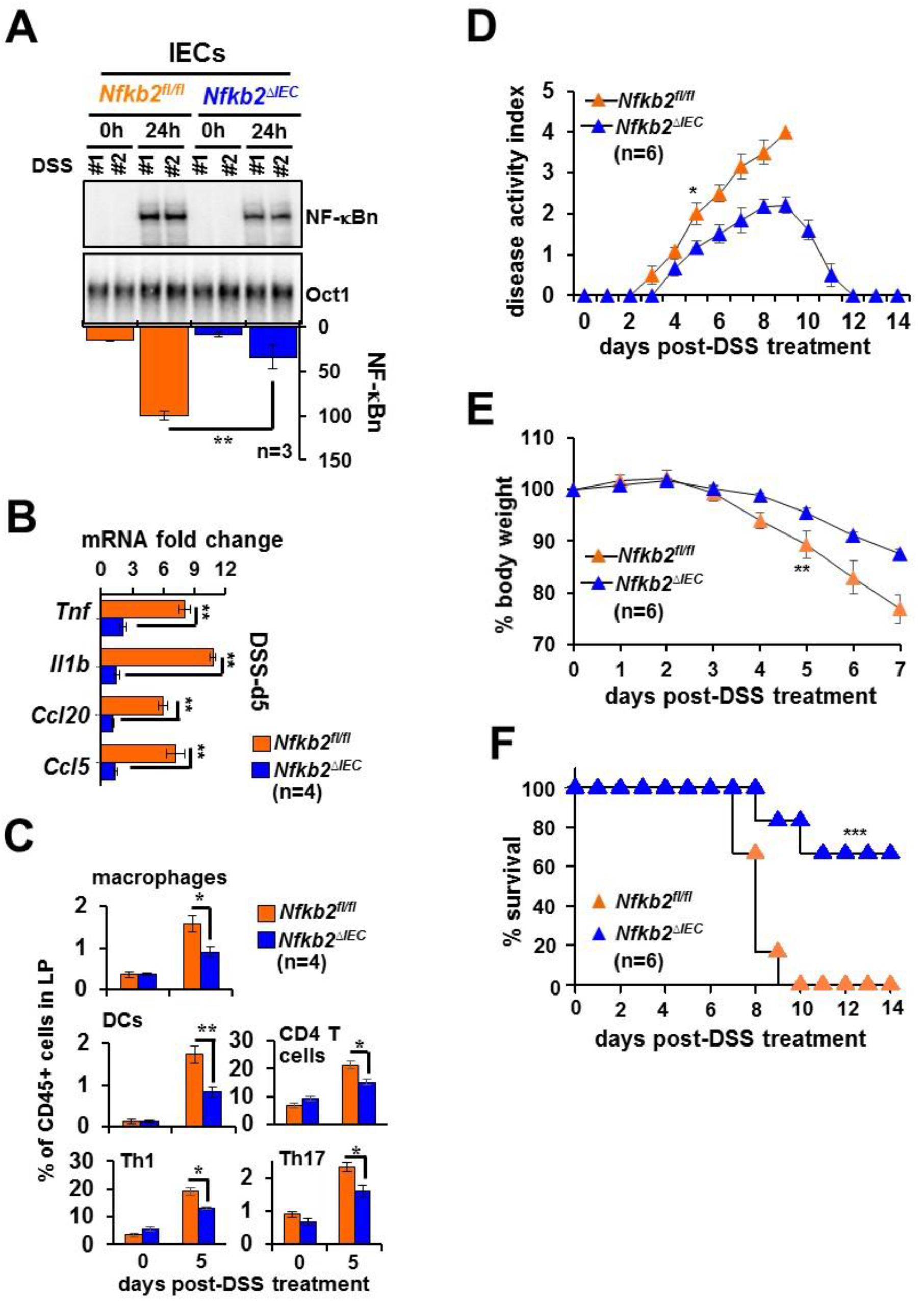
Deficiency of *Nfkb2* in IECs restrains RelA-driven inflammation in the colitogenic gut. **A.** and **B.** IECs isolated from DSS-treated control *Nfkb2^fl/fl^* or *Nfkb2^ΔIEC^* mice were examined for NF-κBn by EMSA **(A)** or pro-inflammatory gene expressions by RT-qPCR **(B)**. **C.** Barplot revealing relative frequencies of the indicated immune cells in the lamina propria of *Nfkb2^fl/fl^* or *Nfkb2^ΔIEC^* mice subjected to DSS treatment. **D.** to **F.** *Nfkb2^ΔIEC^* and *Nfkb2^fl/fl^* mice were subjected to DSS treatment and evaluated for the disease activity **(D)**, bodyweight changes **(E)** and mortality **(F)**. Log-rank (Mantel–Cox) test was used in Figure **4F**. Two-tailed Student’s t-test was performed in other instances. Quantified data represent means ± SEM. \*\*\**P* < 0.001; \*\**P* < 0.01; \**P* < 0.05.

### LTβR-*Nfkb2* signal amplifies canonical NF-κB responses by supplementing latent RelA dimers

We then sought to examine the mechanism linking noncanonical signaling to the pro-inflammatory RelA activity. In their anatomic niche, IECs receive tonic signals through LTβR from innate lymphoid cells expressing the cognate lymphotoxin ligand (Upadhyay and Fu, 2013). To recapitulate this chronic LTβR signaling ex vivo, we first stimulated mouse embryonic fibroblasts (MEFs) for 36h using 0.1μg/ml of an agonistic anti-LTβR antibody (αLTβR), which activates NIK-dependent noncanonical signaling (Banoth et al., 2015). We then activated the canonical pathway in these lymphotoxin-conditioned cells by treating them with microbial-derived LPS in the continuing presence of αLTβR. Our EMSA analyses revealed that chronic LTβR signaling alone induced a minor NF-κBn activity in WT MEFs (Figure 5A). In lymphotoxin-naïve cells, LPS triggered a moderate, RelA-containing NF-κBn activity that peaked at 1h and constituted a weakened late phase. Lymphotoxin conditioning considerably strengthened both the early nRelA peak as well as the late activity induced by LPS. LPS primarily activated RelA:p50 in lymphotoxin-naïve cells (Figure 5B). Lymphotoxin conditioning not only potentiated RelA:p50 activation but also led to a substantial nuclear accumulation of RelA:p52 in response to LPS. As such, *Nfkb2^−/−^* MEFs lack RelA:p52 heterodimers. Importantly, chronic LTβR signaling was ineffective in enhancing even the LPS-induced RelA:p50 activity in *Nfkb2^−/−^* cells (Figure 5A). These studies establish a cell-autonomous role of the LTβR-stimulated noncanonical *Nfkb2* pathway in reinforcing canonical RelA activity.

**Figure 5:**
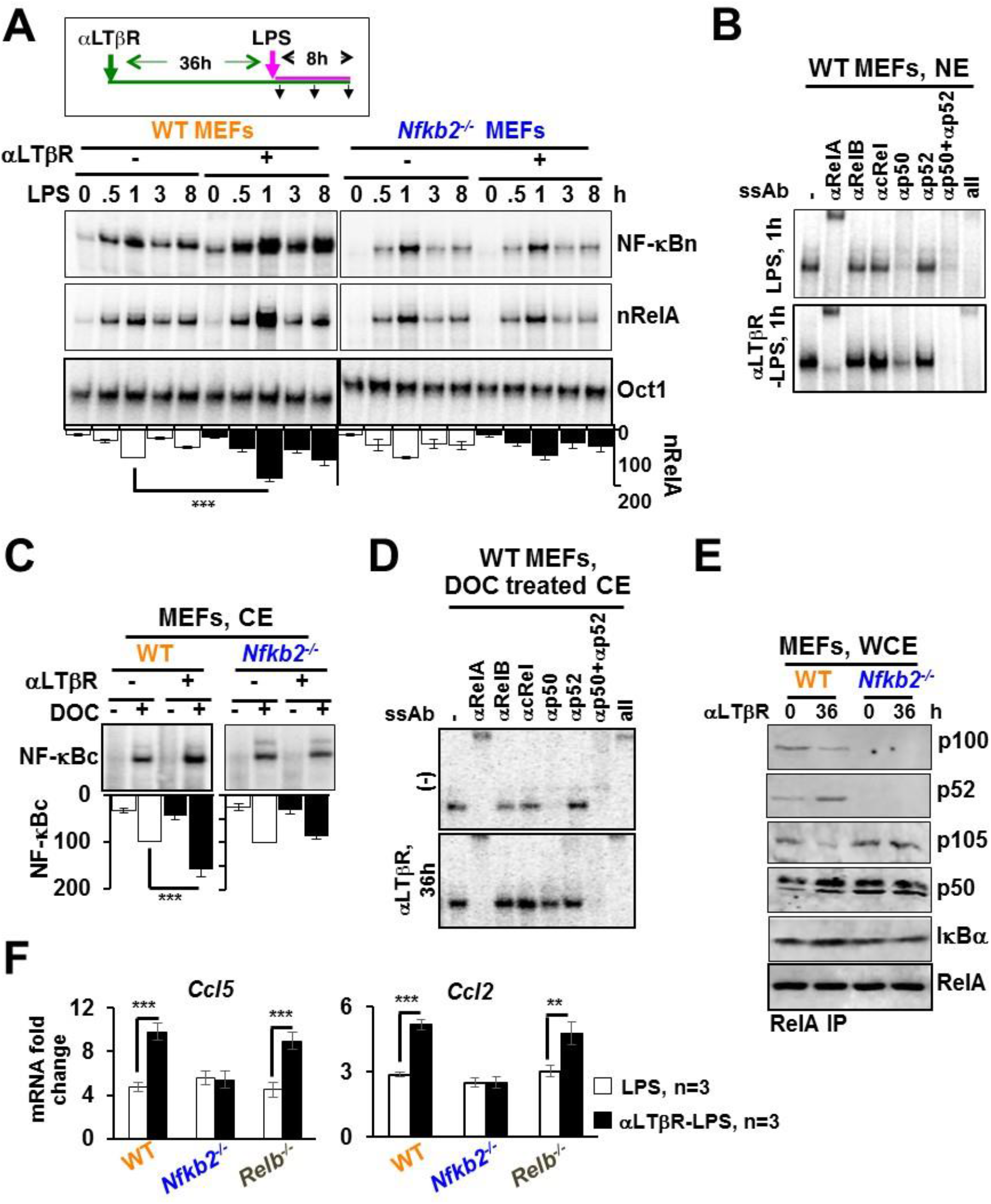
*Nfkb2*-dependent accumulation of latent RelA dimers in LTβR-stimulated MEFs augments canonical NF-κB responses. **A.** EMSA revealing NF-κBn induced in WT and *Nfkb2^−/−^* MEFs in a stimulation time course (top panel). Cells were treated with LPS alone or stimulated using agonistic αLTβR antibody for 36h and then treated with LPS in the continuing presence of αLTβR. Ablating RelB and cRel DNA binding activity using anti-RelB and anti-cRel antibodies, nRelA complexes were examined (middle panel). Oct1 DNA binding (bottom panel) served as a control. The data represent three experimental replicates. **B.** The composition of NF-κBn complexes induced in naïve or LTβR-conditioned MEFs subjected LPS treatment for 1h was characterized by shift-ablation assay. **C.** EMSA revealing the abundance of latent NF-κB dimers in the cytoplasm (NF-κBc) of WT and *Nfkb2^−/−^* MEFs subjected to αLTβR stimulation for 36h. Cytoplasmic extracts were treated with deoxycholate (DOC) before being subjected to EMSA for unmasking latent NF-κB DNA binding activities. The data represent three experimental replicates. **D.** The composition of NF-κBc complexes accumulated in the cytoplasm of MEFs subjected to the indicated treatment regimes was determined by shift-ablation assay. **E.** Immunoblot of RelA coimmunoprecipitates obtained using whole-cell extracts derived from αLTβR-treated MEFs. Data represent three biological replicates. **F.** qRT-PCR analyses revealing pro-inflammatory gene-expressions induced by LPS in naïve or LTβR-conditioned MEFs of the indicated genotypes. Quantified data represent means ± SEM. Two-tailed Student’s t-test was performed. \*\*\**P* < 0.001; \*\**P* < 0.01; \**P* < 0.05.

Canonical signaling entails NEMO-IKK mediated activation of preexisting RelA dimers from the latent, cytoplasmic complexes. It was suggested that NIK might directly regulate the NEMO-IKK activity induced via the canonical pathway (Zarnegar et al., 2008). We found that the low dose of αLTβR used in our experiments, although promoted efficient processing of p100 into p52, did not impact LPS-induced NEMO-IKK activity (Figure S5A and Figure S5B). We then argued that increased availability of latent RelA dimers in the cytoplasm led to hyperactive LPS response in lymphotoxin-conditioned cells. Treatment of cytoplasmic extracts with the detergent deoxycholate (DOC) dissociates RelA dimers from IκBs, unmasking DNA binding activity of the preexisting RelA dimers present in the latent complexes (Baeuerle and Baltimore, 1988). We scored latent NF-κB complexes by analyzing DOC-treated extracts in EMSA. Lymphotoxin conditioning significantly augmented the abundance of latent NF-κB dimers in WT, but not *Nfkb2^−/−^*, MEFs (Figure 5C and Figure S5C). Our shift-ablation assay revealed that the latent NF-κB activity in lymphotoxin-naïve cells consisted of RelA:p50 (Figure 5D). Lymphotoxin conditioning elevated the level of latent RelA:p50 complexes and also accumulated latent RelA:p52 dimer in WT MEFs.

As also reported previously (Yilmaz et al., 2014), LTβR signaling stimulated simultaneous processing of both p105 to p50 and p100 to p52 in our experiments (Figure S5A); although the extent was less remarkable in MEF extracts, signal-responsive p105 processing required the presence of p100 (Figure S5A). Our immunoblot analyses involving RelA co-immunoprecipitates rather clearly revealed that αLTβR treatment of WT MEFs led to an increased association of RelA with p50 and p52 at the expense of RelA binding to the respective p105 and p100 precursors. We also noticed an augmented level of IκBα in the RelA immunoprecipitates derived from LTβR-stimulated WT cells (Figure 5E). Notably, RelA binding to p50 was insensitive to LTβR signaling in *Nfkb2^−/−^* MEFs (Figure 5E). Our analyses suggest that the noncanonical *Nfkb2* pathway targets high-molecular-weight NF-κB precursor complexes to supplement RelA:p50 and RelA:p52 heterodimers, which are then sequestered by IκBα in the latent NF-κB complexes.

Notably, RelA:p50 and RelA:p52 was shown to exhibit overlap in relation to pro-inflammatory gene expressions (Banoth et al., 2015; Hoffmann and Leung, 2003). Accordingly, lymphotoxin conditioning amplified LPS-induced expressions of generic RelA target chemokine genes encoding Ccl5 and Ccl2 in WT, but not *Nfkb2^−/−^*, cells (Figure 5F). LTβR signaling also stimulates nuclear RelB activity. Lymphotoxin-mediated enhancement of LPS-induced gene expressions in *Relb^−/−^* MEFs asserted that the observed gene effects were independent of RelB. We infer that tonic LTβR-*Nfkb2* signaling increases the abundance of latent RelA heterodimers leading to hyperactive canonical NF-κB response, which amplifies TLR-induced expressions of RelA-target pro-inflammatory genes.

### LTβR-*Nfkb2* signal in the intestinal niche instructs the homeostasis of latent RelA dimers in IECs

Next, we asked if the *Nfkb2* pathway modulated latent NF-κB complexes also in the intestinal niche. Immunoblot analyses of extracts from IECs revealed a muted processing of p105 to p50 in *Nfkb2^−/−^* compared to WT mice (Figure 6A). The reduced cellular abundance of p50 in *Nfkb2^−/−^* mice led to decreased RelA binding to p50 (Figure 6B). On the other hand, treatment of WT mice with LTβR-Ig prevented p100 processing to p52 and also diminished p50 production from p105. LTβR-Ig treatment of WT mice lessened the association of RelA with both p50 and p52. The latent NF-κB activity in IECs consisted of mostly RelA:p50, and a moderate amount of RelA:p52, in WT mice (Figure 6C and Figure 6D). Global (Figure 6C) or IEC-specific (Figure 6E) *Nfkb2* deficiency or LTβR-Ig treatment (Figure 6F) substantially reduced the abundance of latent RelA dimers in IECs. Together, tonic LTβR-*Nfkb2* signaling determines the homeostasis of latent RelA dimers in IECs by promoting simultaneous production of RelA-interacting partners p50 and p52 from their respective precursors.

**Figure 6:**
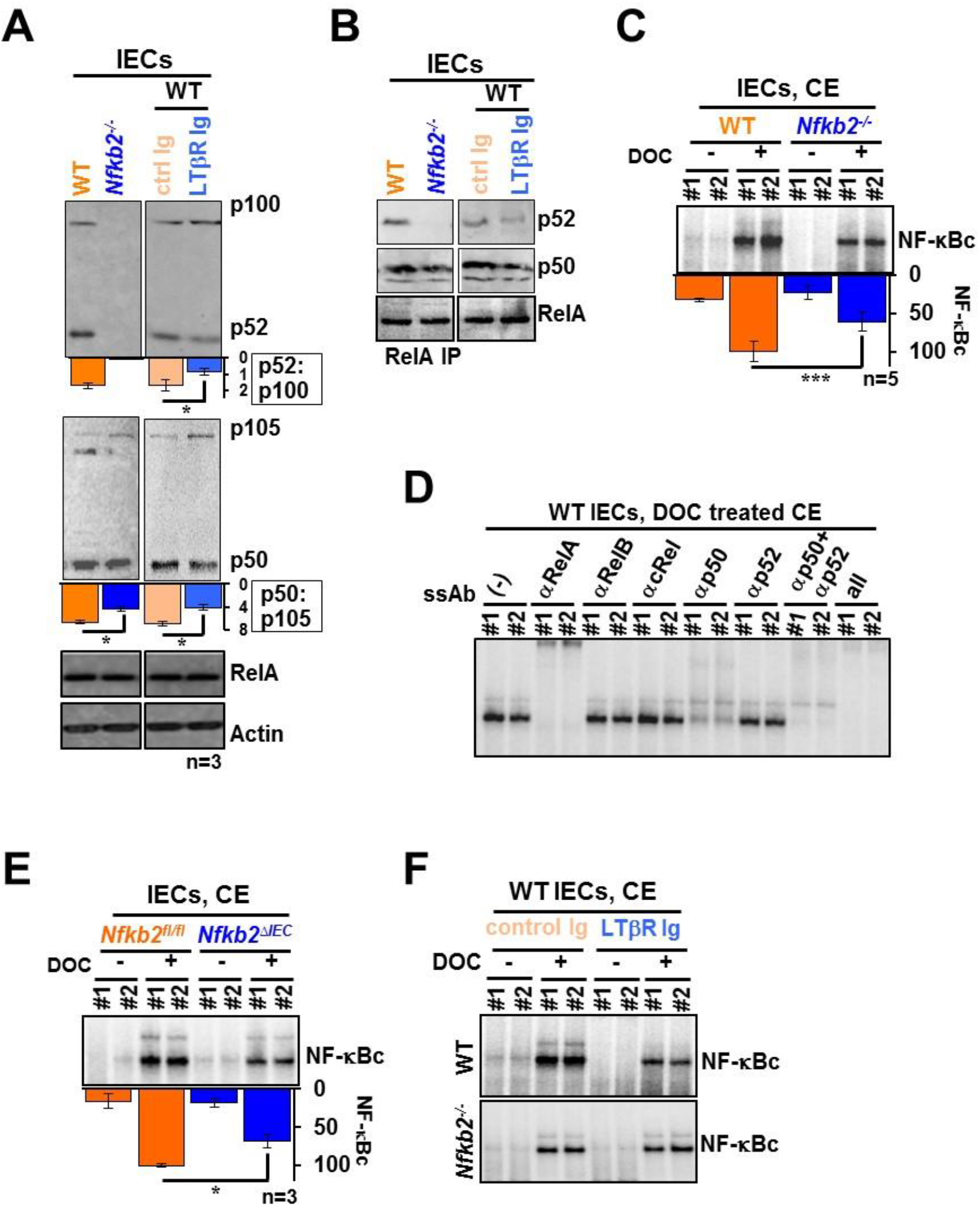
LTβR-*Nfkb2* signaling regulates the abundance of latent RelA dimers in IECs. **A.** and **B.** Immunoblot of whole-cell extracts **(A)** or RelA coimmunoprecipitates obtained using whole-cell extracts **(B)** derived from IECs. IECs were obtained from WT or *Nfkb2^−/−^* mice or WT mice administered with either control-Ig or LTβR-Ig 24h prior to tissue collection. Quantified signals corresponding to p52:p100 or p50 to p50:p105 has been indicated below the respective lanes. **C.** and **E.** and **F.** EMSA revealing the abundance of latent NF-κBc dimers present in the cytoplasm of IECs derived from the indicated mice. Cytoplasmic extracts were treated with DOC for unmasking latent NF-κB DNA binding activity. Data represent five **(C)** or **(E** and **F)** three biological replicates. **D.** The composition of latent NF-κBc dimers present in IECs derived from WT mice was determined by shift-ablation assay. Quantified data represent means ± SEM. Two-tailed Student’s t-test was performed. \*\*\**P* < 0.001; \**P* < 0.05.

### Increased availability of p50 and p52 connects elevated noncanonical NF-κB signaling to heightened nRelA activity in human IBD

Because human IBD was associated with increased p100 processing to p52, we asked if IBD patients also exhibited increased p50 production from p105. Immunoblot analyses of extracts derived colonic epithelial biopsies identified a total of twenty-three IBD patients with heightened p105 processing, as assessed from a one-and-a-half-fold or more increase in the p50:p105 ratio compared to the median value in the control cohort (Figure 7A and Figure S1D). There was an overall 1.7 fold increase in the median p50:p105 value in IBD patients. In an upset plot, we could further capture that the fraction of patients displaying elevated ratios of both p50:p105 and p52:p100 was substantially higher than those with augmented ratios of either p50:p105 or p52:p100 (Figure 7B). A high odds ratio for a simultaneous increase in the p50:p105 and p52:p100 values indicated interdependent processing of p105 and p100 in IBD. Finally, odds ratio measurements established that heightened nRelA activity in IBD was closely linked with increased processing of both the NF-κB precursors (Figure 7C). These studies ascribe increased availability of the RelA dimerization partners p50 and p52, generated through interdependent processing of NF-κB precursors, to the heightened RelA activity in human IBD.

**Figure 7:**
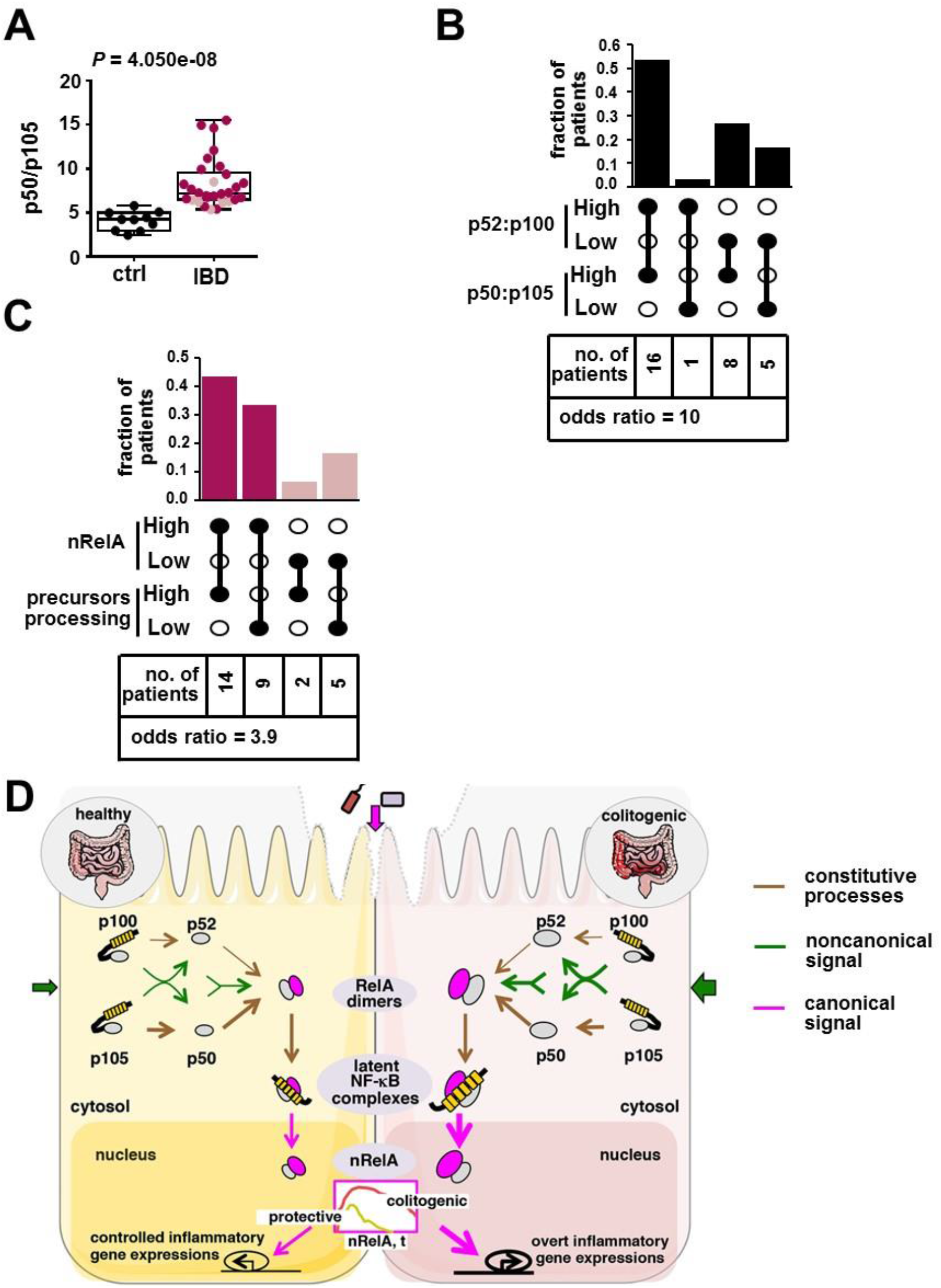
A link between elevated processing of p100 as well as p105 and heightened nRelA activity in human IBD. **A.** The abundance of p50 in relation to p105 in colon biopsies from controls and IBD patients was quantified involving immunoblot analyses (see Figure S1D) and presented as a dot plot. Statistical significance was determined by Welch’s t-test. **B.** Upset plot showing the fraction of IBD patients with elevated processing of both p100 and p105. Patients with p52:p100 ratio above 1.5 fold of the median of the control cohort were termed as “p52:p100 High”; they were otherwise considered as “p52:p100 Low”. Similarly, patients were catalogued as either p50:p105 High or p50:p105 Low. The number of patients in each category has been indicated. The interdependence of p100 and p105 processing was determined from the odds ratio of a patient to have high p50:p105 ratio if p52:p100 ratio was elevated (90% CI, 1.44 to 69.41). **C.** Upset plot revealing the fraction of IBD patients with heightened nRelA activity as well as increased processing of both the NF-κB precursors p100 and p105. The interdependence was determined from the odds ratio (90% CI, 0.83 to 18.24). **D.** A mechanistic model explaining proposed role of the noncanonical *Nfkb2* pathway in amplifying canonical RelA activity that causes aberrant inflammation in the colitogenic gut. In this cartoon, brown, magenta and green lines represent constitutive, canonical signal-responsive and noncanonical signal-responsive cellular processes, respectively. The weight of the lines indicates the strength of the respective biochemical reactions.

Taken together, we put forward a mechanistic model explaining aberrant intestinal inflammation (Figure 7D). In this model, noncanonical *Nfkb2* signaling in IECs supplemented latent RelA heterodimers aggravating canonical NF-κB response in the colitogenic gut. We suggest that latent RelA dimer homeostasis connects the noncanonical NF-κB pathway to RelA-driven inflammatory pathologies.

## Discussion

Despite the well-established role of RelA in fueling aberrant intestinal inflammation, the molecular mechanism underlying inordinate RelA activation in the colitogenic gut remains unclear. As such, the canonical NF-κB pathway drives controlled nuclear activity of RelA. Our investigation revealed not only the triggering of canonical signaling but also the frequent engagement of the noncanonical NF-κB pathway in human IBD. In a mouse model, we ascertained that tonic, noncanonical LTβR-*Nfkb2* signaling amplified epithelial RelA activity induced during the initiation of experimental colitis. This *Nfkb2*-mediated regulation escalated RelA-driven pro-inflammatory gene response in IECs, exacerbating the infiltration of inflammatory cells and colon pathologies. Our analyses involving cultured cells confirmed that noncanonical LTβR-*Nfkb2* signaling intensified canonical response to TLRs in a cell-autonomous mechanism. Although multifactorial human IBD cannot be fully recapitulated in animal models, our finding suggests that impinging noncanonical signaling aggravates inflammatory, canonical RelA activity in the colitogenic gut of IBD patients. Unlike basally elevated noncanonical signaling in the mouse colon, however, p100 processing was rather timid in IBD-free human subjects. While this difference warrants further examination, future studies ought to determine if the onset of human IBD is indeed preceded by strengthening *Nfkb2* signaling or engagement of the noncanonical pathway coincides with the disease progression. What provides noncanonical signals particularly in human IBD also remains unclear; we suggest that investigating the dynamic engagement of immune cells bearing ligands for LTβR may offer important insights.

Previous mouse studies revealed a rather complex role of the canonical pathway in the colon (Wullaert et al., 2011; Zaidi and Wine, 2018). Genetic deficiency of the inhibitory IκBα in IECs caused spontaneous intestinal inflammation (Mikuda et al., 2020). However, a complete lack of canonical signaling in IEC-specific knockouts of RelA or IKK2 sensitized those mice to experimental colitis despite reduced inflammatory gene activation (Eckmann et al., 2008; Steinbrecher et al., 2008). In addition to immune genes, RelA also activates the expression of antiapoptotic factors. It was found that increased apoptosis of IECs in the absence of functional canonical signaling overwhelms mucosal healing. Targeting pathway components using antisense oligo or peptide inhibitors achieved partial inhibition of canonical signaling in mice subjected to colitogenic insults (Neurath et al., 1996; Shibata et al., 2007). Interestingly, such incomplete pathway inhibition effectively alleviated gut inflammation while circumventing cellular apoptosis in the intestinal niche. An absence of *Nfkb2* diminished, but not completely abrogated, canonical RelA signaling in IECs in our study. Accordingly, *Nfkb2* deficiency restrained intestinal inflammation without exacerbating cell death in the colitogenic gut.

Investigating the colitogenic function of the noncanonical pathway in gene knockout mice yielded confounding outcomes. Increased intestinal inflammation observed in *Nlrp12^−/−^* mice was attributed to the elevated level of NIK and p52 (Allen et al., 2012). We have also earlier reported that *Nfkb2* signaling reinforces protective inflammatory responses against gut pathogens (Banoth et al., 2015). Our current study revealed that *Nfkb2*-mediated amplification of canonical signaling in IECs rather conferred vulnerability in mice enduring chemically-induced, pervasive colon damage. While our finding was consistent with those published by Burkitt et al. (2015), we further established the role of LTβR in triggering this *Nfkb2* pathway in IECs. However, LTβR-Ig was unable to ameliorate DSS-induced colitis (Jungbeck et al., 2008). More so, IEC-specific ablation of either LTβR or NIK sensitized mice to DSS (Macho-Fernandez et al., 2015; Ramakrishnan et al., 2019). While these studies did not causally link NF-κB precursor processing to intestinal inflammation, NIK was found to be essential for maintaining M cells, which produced gut protective IL-17A and IgA (Ramakrishnan et al., 2019). We reconcile that LTβR and NIK limit experimental colitis involving *Nfkb2*-independent M cell functions, while LTβR-driven *Nfkb2* signaling in less-specialized IECs aggravates colon inflammation. In support of our model, NIK’s overexpression in IECs upregulated the colonic expression of pro-inflammatory cytokines in mice (Ramakrishnan et al., 2019), and human IBD was associated with an augmented colonic abundance of LIGHT, an LTβR ligand (Cohavy et al., 2005).

The amplitude of canonical RelA response depends on both the magnitude of signal-induced NEMO-IKK activity as well as the abundance of IκB-bound latent dimers, which are acted upon by NEMO-IKK. While NEMO-IKK activation has been extensively studied in the context of inflammation, molecular processes regulating the generation of latent NF-κB dimers remain less explored. It was demonstrated that by preferentially stabilizing RelA homodimers in resting cells, IκBβ ensured their availability for activation via the canonical pathway (Tsui et al., 2015). Our study revealed that noncanonical signaling tuned the homeostasis of latent RelA heterodimers. Scheidereit’s group earlier reported that LTβR stimulates simultaneous processing of p100 and p105 and that LTβR-responsive p105 processing to p50 requires p100 (Yilmaz et al., 2014). We found that the noncanonical pathway accumulated both RelA:p50 and RelA:p52 involving this interdependent processing mechanism. Sequestration of these RelA dimers by IκBs in deoxycholate-sensitive latent complexes provided for heightened canonical response in lymphotoxin-conditioned cells. The absence of *Nfkb2* diminished LTβR-responsive, but not constitutive, RelA:p50 generation. We propose that the previously-described high-molecular-weight complexes comprising RelA, p105, and p100 (Savinova et al., 2009) produced latent RelA dimers in response to LTβR signal and that a lack of p100 prevented proteolytic machinery from recruiting to these complexes. Nonetheless, our data suggested that interdependent processing of p105 and p100 modulated intestinal inflammation in colitogenic mice and in human patients. Despite impinging noncanonical signaling in the gut, however, we could detect only a minor nuclear RelB DNA binding activity in IECs. We speculate that an IEC-intrinsic mechanism may either prevented RelB activation or abrogated nuclear RelB activity in the inflamed colon.

Nonetheless, we present evidence that the convergence of seemingly harmless, tonic signals with inflammatory signals may provoke pathological RelA activity. Within the intestinal niche, epithelial cells receive noncanonical NF-κB signals, which inflate the repertoire of latent RelA dimers; mucosal damage provides for the inflammatory signal via the canonical NF-κB pathway that then fuels aberrant RelA activity in the gut. Direct therapeutic targeting of the canonical pathway precipitates undesired side effects because of the engagement of canonical signaling in a myriad of physiological processes. In this context, the noncanonical pathway offers an attractive therapeutic option. Notably, small molecule inhibitors of NIK showed promising results in mitigating experimental lupus in mice (Brightbill et al., 2018). We argue that targeting NIK-mediated regulations high-molecular-weight precursor complexes may tune the NF-κB dimer homeostasis alleviating RelA-driven inflammatory pathologies, including those associated with IBD. In sum, our study emphasizes that signal-crossregulatory mechanisms may provide for specific therapeutic targets in human ailments.

## STAR◽METHODS

### KEY RESOURCES TABLE

2 KEY RESOURCES TABLE
| Reagent | Source | Identifier |
| --- | --- | --- |
| **Antibodies** |  |  |
| RelA Rabbit Polyclonal IgG | Santa Cruz Biotechnology | sc-372 |
| RelB Rabbit Polyclonal IgG | Santa Cruz Biotechnology | sc-226 |
| cRel Rabbit Polyclonal IgG | Santa Cruz Biotechnology | sc-71 |
| IκBα Rabbit Polyclonal IgG | Santa Cruz Biotechnology | sc-371 |
| *Nfkb2* p100/p52 Antibody | Cell Signaling Technology | 4882 |
| Anti-p50 Antibody | BioBharati LifeSciences | BB-AB0080 |
| Anti-NEMO Antibody | BioBharati LifeSciences | BB-AB0035 |
| IKK1 Rabbit Polyclonal IgG | Santa Cruz Biotechnology | sc-7184 |
| β-Actin Goat Polyclonal IgG | Santa Cruz Biotechnology | sc-1615 |
| Goat anti-Rabbit IgG-HRP Secondary Antibody | Santa Cruz Biotechnology | sc-2004 |
| Donkey anti-Goat IgG-HRP Secondary Antibody | Santa Cruz Biotechnology | sc-2020 |
| Goat anti-Rabbit IgG-Cy5 Secondary Antibody | GE-Healthcare | 29038278 |
| Rabbit Trueblot Anti-Rabbit IgG HRP | Rockland | 18-8816-33 |
| Anti-mouse CD45.2 APC Antibody | eBiosciences | 17-0454-82 |
| Anti-mouse CD45.1 PE Antibody | eBiosciences | 12-0453-82 |
| Anti-mouse EpCAM PE Antibody | eBiosciences | 12-5791-82 |
| Anti-mouse CD90.2 FITC Antibody | Biolegend | 105306 |
| Anti-human/mouse CD45R PE Antibody | eBiosciences | 12-0452-82 |
| Anti-mouse CD19 FITC Antibody | eBiosciences | 11-0191-82 |
| Anti-mouse NK1.1 FITC Antibody | eBiosciences | 11-5941-82 |
| Anti-mouse CD11b brilliant violent 510 Antibody | Biolegend | 101245 |
| Anti-mouse CD11c PerCP Antibody | Biolegend | 117236 |
| Anti-mouse Lys-6G FITC Antibody | eBiosciences | 11-9668-82 |
| Anti-mouse F4/80 eFluor 450 Antibody | eBiosciences | 48-0251-82 |
| Anti-mouse MHCII PE-Cy7 Antibody | eBiosciences | 25-5321-82 |
| Anti-mouse CD64 PerCP-e710 Antibody | eBiosciences | 46-0641-82 |
| Anti-mouse CD4 PE-Cy7 Antibody | eBiosciences | 25-0041-82 |
| Anti-mouse CD25 eFluor 450 Antibody | eBiosciences | 48-0251-82 |
| Anti-mouse IFNγ APC-Cy7 Antibody | Biolegend | 505850 |
| Anti-mouse/rat IL17A PerCP-Cy5.5 Antibody | eBiosciences | 45-7177-82 |
| Anti-mouse/rat Foxp3 FITC Antibody | eBiosciences | 11-5773-82 |
| Anti-mouse Siglec F APC-Cy7 Antibody | BD Bioscience | 565527 |
| **Chemicals and Peptides** |  |  |
| DSS (36,000-50,000 MW) | MP Biomedicals | 160110 |
| FITC-Dextran (MW 3000 - 5000) | Sigma-Aldrich | 60842-46-8 |
| Agonistic LTβR antibody (αLTβR), AFH6 | Biogen Inc., USA | AFH6 |
| Antagonistic LTβR-Ig Antibody | Biogen Inc., USA | N/A |
| Control MOPC21 Antibody | Biogen Inc., USA | N/A |
| Collagenase, Type 4 | Worthington | LS004188 |
| DNase I | Worthington | LS002139 |
| LPS from E.coli serotype O55:B5 | Enzo Life Sciences | ALX-581-013-L002 |
| Recombinant GST-IkBa (1-54) protein | BioBharati Life Sciences | Customized |
| **Commercial Kits** |  |  |
| RNeasy Mini Kit (250) | Qiagen | 74106 |
| Primescript 1<sup>st</sup> strand cDNA synthesis kit | Takara Bio | 6110B |
| Foxp3/Transcription Factor Staining Buffer Set | eBioscience | 00-5523-00 |
| **Mice models** |  |  |
| C57BL/6 WT mice | available at the Small Animal Facility in NII | N/A |
| C57BL/6 SJL mice | Jackson Laboratories | 002014 |
| *Nfkb2*<sup>-/-</sup> mice | available at the Small Animal Facility in NII. | N/A |
| *Nfkb2*<sup>fl/fl</sup> mice | Jackson Laboratories | 028720 |
| Villin-Cre mice | gift from Prof. Florian Greten, Georgspeyer-Haus Institute of Tumor Biology and Experimental Therapy, Germany. | N/A |
| *Nfkb2*<sup>ΔIEC</sup> mice | generated by crossbreeding *Nfkb2*<sup>fl/fl</sup> mice with Villin-Cre mice. | N/A |

#### Primers for qRT-PCR

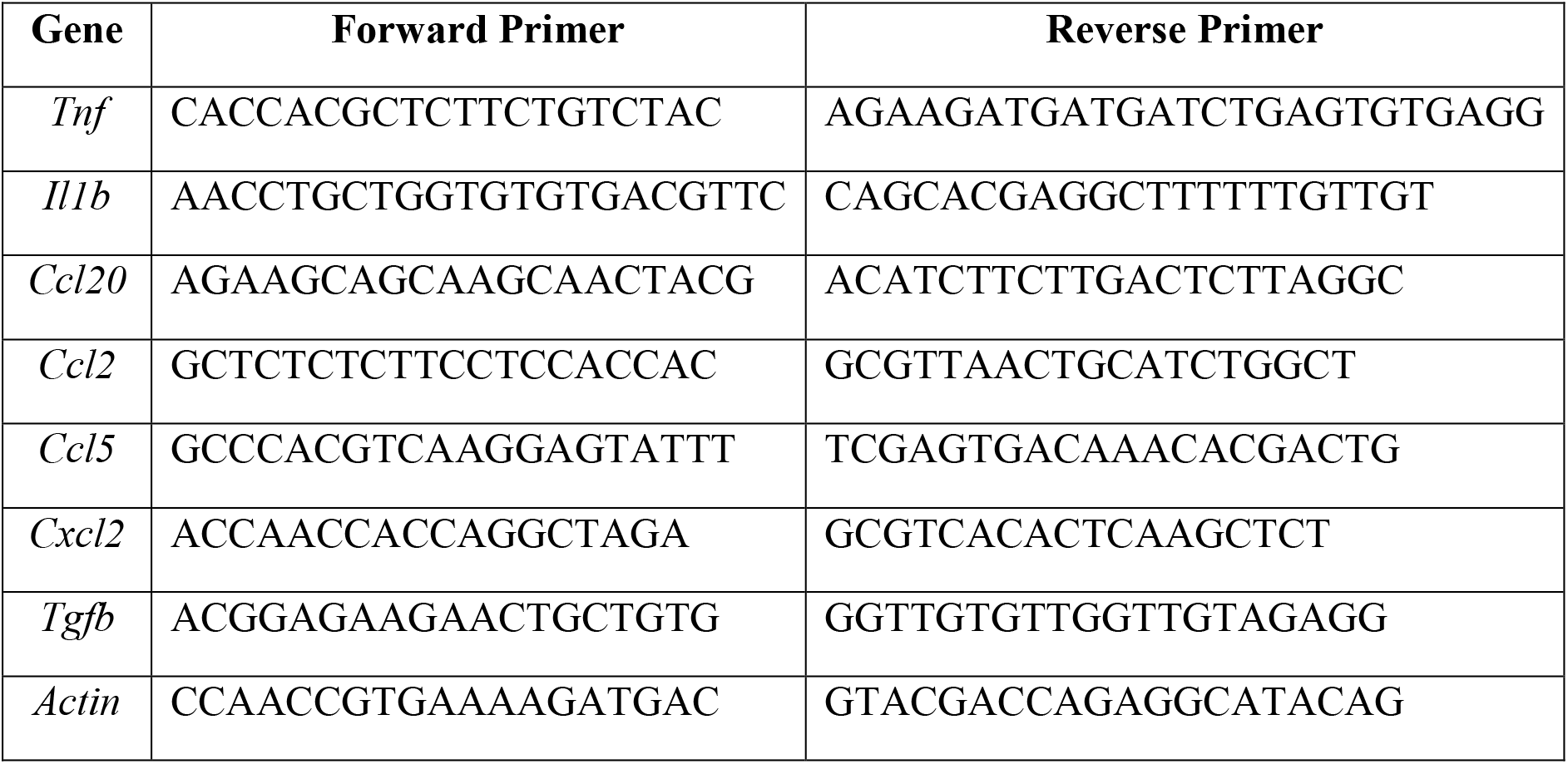

##### Patients and collection of colon biopsies

Patients older than 18 years were registered at the All India Institute of Medical Sciences, New Delhi, India, and diagnosed based on European Crohn’s and Colitis Organization (ECCO) guidelines. We examined IBD patients suffering from ulcerative colitis. Colonic epithelial biopsies were derived from the inflamed region of the rectum and subjected to biochemical analyses. As controls, we utilized samples from IBD-free individuals suffering from hemorrhoids. Experimental procedures were approved by the institutional human ethics committees of the All India Institute for Medical Sciences (Protocol No. IHEC-667/07-12-2018) and the National Institute of Immunology (Protocol No. IHEC#106/18), and informed consent was obtained from patients.

##### Animal use

All mouse strains were housed at the National Institute of Immunology (NII) and used adhering to the institutional guidelines (Approval number – IAEC 400/15). *Nfkb2^fl/fl^* mice (stock no. #028720, C57BL/6) were obtained from the Jackson Laboratory and Villin-Cre mice were a generous gift from Dr. Florian Greten, Georgspeyer-Haus Institute of Tumor Biology and Experimental Therapy, Germany. Villin-Cre mice were crossed with *Nfkb2^fl/fl^* mice to generate *Nfkb2^Δ^*^CEI^ mice, which lacked *Nfkb2* functions specifically in IECs. *Nfkb2^Δ^*^CEI^ and control *Nfkb2^fl/fl^* mice used in our experiments were in the mixed background.

##### Induction and assessment of colitis in mice

As described earlier (Kiesler et al., 2015), 7-9 weeks old male mice of the indicated genotypes were administered with 2.5% of DSS in drinking water for seven days. Subsequently, mortality, bodyweight, and disease activity were assessed for fourteen days from the onset of DSS treatment. All experiments were performed using littermate mice cohoused for a week prior to the initiation of the experiments. The disease activity index was estimated based on stool consistency and rectal bleeding. The score was assigned as follows – 0 points were given for well-formed pellets, 1 point for pasty and semi-formed stool, 2 points for liquid stool, 3 points for bloody smear along with stool, and 4 points were assigned for bloody fluid/mortality. Mice with more than 30% loss of bodyweight were considered moribund (Franco et al., 2012), and euthanized. Data from moribund mice have not been included in the survival plot. For specific experiments, mice were euthanized at the indicated days post-onset of DSS treatment, and colon tissues were collected. 200 μg of antagonistic LTβR-Ig was injected via intraperitoneal route into mice as described earlier (Banoth et al., 2015), and IECs were collected after another 24 h. As a control, MOPC21 antibody was used.

##### Colon length and histological studies

At day seven of DSS treatment, mice with the indicated genotypes were euthanized, and the entire colon was excised. The colon length was measured as the length from the rectum to the caecum. Subsequently, distal colons were washed with PBS, fixed in 10% formalin and embedded in paraffin. 5μm thick tissue sections were generated from the inflamed region and stained with hematoxylin and eosin (H&E). Alternately, sections were stained using H&E as well Alcian Blue for revealing mucin content. Images were captured using Image-Pro6 software on an Olympus inverted microscope under a 20X objective lens. The severity of colitis was assessed by epithelial damage and infiltration of inflammatory immune cells in the submucosa of the colon. For assessing intestinal permeability, FITC-dextran were gavaged to DSS-treated mice 6h prior to analysis of FITC fluorescence in serum.

##### Generation of bone marrow chimeras

Eight weeks old CD45.2^+^ or CD45.1^+^ mice were irradiated with 10 Gy of radiation and then administered retro-orbitally with 1-2 ×10^7^ bone marrow cells derived from tibia and femur of the donor CD45.1^+^ or CD45.2^+^ mice, respectively. After four weeks of transfer, chimerism was examined involving flow cytometric analyses of CD45.1 and CD45.2 markers on peripheral lymphocytes in the recipient mice. Subsequently, mice were treated with DSS.

##### Isolation of IECs from mouse colons

Colons were surgically removed from the indicated mice subsequent to euthanization, longitudinally cut open, washed with PBS, and then cut into 2-5 mm small pieces. These pieces were rocked on a shaker platform submerging in HBSS containing 30mM EDTA that led to the detachment of IECs. IECs were collected by centrifugation and subjected to biochemical analyses.

##### Isolation of lamina propria cells from mouse colons

Colons harvested from the indicated mice were first longitudinally cut open, and then cut into small pieces. These small pieces were rocked on a shaker platform submerging in HBSS containing 1mM EDTA, 1mM DTT and 10% FBS. Subsequently, pieces were washed gently and then subjected to enzymatic digestion for 60 min in HBSS containing 0.5 mg/ml collagenase IV (Worthington), 0.05 mg/ml DNase I (Worthington) and 10% FBS. The supernatant was filtered through a nylon cell strainer with 70μm pore size (Corning). Cells from the filtrate were collected, stained with fluorochrome-conjugated antibodies and analyzed by flow cytometry.

##### Flow cytometric analyses

Fluorochrome conjugated antibodies against mouse CD45.1 (A20), CD45.2 (104), CD90.2 (30-H12), B220 (RA3 6B2), CD19 (MB19), NK1.1 (PK136), Ly6G (1A8), SiglecF(E50-2440), MHCII (IA/IE), CD64 (X53-5/7.1), CD4 (GK1.5), CD11c (N418), EpCAM (G8.8) were purchased from Ebioscience, BD biosciences or Biolegend. Flow cytometry was performed using FACS Verse flow cytometer (BD Biosciences). Data was analyzed using FlowJo v9.5 software (Treestar, Ashland and OR). Briefly, neutrophils were identified as Gr1+SiglecF-cells; macrophages and local dendritic cells as CD11b+F4/80+ and CD11c+F4/80-, respectively; and Th1, Th17 and Treg cells were recognized as CD4+IFNγ+, CD4+IL17A+ and CD4+CD25+FoxP3+ cells, respectively.

##### Gene expression studies

Total RNA was isolated using RNeasy Mini Kit (Qiagen, Germany) from IECs isolated from DSS treated WT and *Nfkb2^−/−^* mice. cDNA was synthesized using Reverse Transcriptase (Takara Bio Inc.) and oligo dT primers. qRT-PCR were performed using Power SYBR Green PCR master mix (Invitrogen) in ABI 7500 FAST instrument. Relative mRNA levels were determined using the 2^−ΔΔCt^ methods. Actin was used as a control. Total RNA isolated from IECs was further subjected to the RNA-seq analysis at the core facility of the Singapore Immunology Network. IECs were obtained from WT and *Nfkb2^−/−^* mice, either left untreated or treated with DSS (n=3). If the cumulative read count of a transcript estimated from a total of twelve experimental sets was less than five hundred, the corresponding gene was excluded from analyses. Ensemble IDs lacking assigned gene names were also excluded. Accordingly, we arrive onto a list of 8,199 genes from 46,517 entries in our dataset (available in GEO, GSE148577). The average read counts of these genes in various experimental sets were determined. Subsequently, fold changes in the mRNA levels upon DSS treatment were determined using DEseq2 package from Bioconductor (Love et al., 2014). The difference in the fold change values between WT and *Nfkb2^−/−^* mice was calculated for 8,199 genes, and genes were then ranked in descending order of this fold change difference. This ranked gene list was then subjected to the gene set enrichment analysis involving fgsea package using a previously described list of RelA-target genes (Banoth et al., 2015).

##### *Ex vivo* cell stimulations

WT or *Nfkb2^−/−^* MEFs, obtained from day 14.5 embryos, were treated with 0.1 μg/ml of αLTβR or 1 μg/ml of LPS for the indicated times. Alternately, MEFs were stimulated with αLTβR for 36h and then lymphotoxin-conditioned cells were additionally treated LPS in the continuing presence of αLTβR.

##### Biochemical studies

###### Cell extract preparation

Methods for preparing nuclear, cytoplasmic, and whole-cell extracts from MEFs and mouse IECs have been described elsewhere (Banoth et al., 2015). Adapting from a previously published protocol (Andresen et al., 2005), we optimized a nucleo-cytoplasmic fractionation method for colonic epithelial biopsies derived from human subjects. Briefly, colon biopsies were homogenized and suspended in hypotonic cytoplasmic extraction buffer (10mM HEPES-KOH pH 7.9, 10mM KCl, 1.5mM MgCl2, 1mM EDTA, 0.5% NP40, 5% sucrose, 0.75mM spermidine, 0.15mM spermine, 1mM DTT, 1mM PMSF). Samples were centrifuged at 8,000g, cytoplasmic extracts were isolated and nuclear pellets were resuspended in nuclear extraction buffer (250 mM Tris-HCl pH 7.5, 60mM KCl, 1 mM EDTA, 0.5 mM DTT, 1 mM PMSF). Nuclear pellets were then subjected to three freeze-thaw cycles involving dry ice and 37^0^C water bath, and nuclear extracts were isolated subsequent to centrifugation. Total protein was quantified using Lowry assay.

###### EMSA

Detailed protocols for EMSA and shift-ablation assay have been described earlier (Banoth et al., 2015). Briefly, 5-10 μg of nuclear lysate, obtained from either mouse or human cells, was incubated with a ^32^P labeled DNA probe derived from HIV long terminal repeats containing kappaB sites. Samples were resolved on a 6% native gel, and gel was exposed to a storage phosphor screen. Gel images were acquired using Typhoon 9400 Variable Mode Imager and NF-κB signal intensities were quantified in ImageQuant 5.2. For shift-ablation assay, nuclear extracts were first incubated with antibodies against individual NF-κB subunits, and then DNA probe was added. The contribution of a given NF-κB subunit in nuclear DNA binding was then interpreted from the reduction of specific NF-κB-DNA complexes in EMSA gel. In RelA-EMSA, nuclear extracts were preincubated with anti-RelB as well as anti-cRel antibodies that ablated their respective DNA-binding activities, allowing for quantification of RelA DNA binding activity. For assessing latent NF-κB dimers, 2 μg of cytoplasmic extract was treated with 0.8% DOC for 30 min and then analyzed in EMSA subsequent to incubating to DNA probe (Basak et al., 2007).

###### NEMO-IKK kinase assay

As described earlier (Banoth et al., 2015), NEMO coimmunoprecipitates obtained from cytoplasmic extracts were incubated with γ^32^P-ATP and recombinant GST-IκBα. Resultant reaction mixtures were resolved by SDS-PAGE. Gel images were acquired using PhosphorImager; the NEMO-IKK activity was assessed from the extent of phosphorylation of IκBα.

###### Immunoblot analyses

For immunoblot analyses (Banoth et al., 2015), whole-cell extract was prepared in SDS-RIPA buffer (50 mM Tris-HCl pH 7.5, 150 mM NaCl, 1 mM EDTA, 0.1% SDS, 1% Triton-X 100, 1 mM DTT, 1 mM PMSF), and resolved by SDS-PAGE. Subsequently, proteins were transferred onto a PVDF membrane, and immunoblotting was performed using indicated antibodies. In particular, we used Cy5 conjugated secondary antibody for analyzing human colon biopsies. Gel images were acquired using Typhoon Variable Mode Imager and band intensities were quantified in ImageQuant.

###### Immunoprecipitation studies

The whole-cell lysate was prepared from ~ 10^6^ cells in a buffer containing 20 mM of Tris-HCl pH 7.5, 20% of glycerol, 0.5% of NP-40, and a final 150 mM of NaCl (Banoth et al., 2015). Immunoprecipitation was performed by adding ~ 2μgm of anti-RelA antibody to the whole-cell extract. For immunoblot analyses of RelA-coimmunoprecipitates, Trueblot secondary antibody was used.

##### Statistical analysis

Error bars are shown as SEM of 4-8 mice in animal studies and as SEM of 3-5 replicates in biochemical experiments. Quantified data are means ± SEM. Unless otherwise mentioned, paired two-tailed Student’s t-test was used for calculating statistical significance in dataset involving mouse or derived cells. Human data were subjected to Welch’s unpaired t-test.

## Data and Software Availability

The MIAME version of the RNA seq dataset is available on NCBI Gene Expression Omnibus (accession number GSE148577).

## Supplemental Information

Supplemental Information consists of five supplementary figures.

## Acknowledgement

We sincerely thank Biogen Inc. for providing αLTβR and LTβR-Ig antibodies, and Prof. F. Greten for the generous gift of Villin-Cre mice. We thank V. Kumar for technical help; B. Lee for help with RNA-seq analyses; and Dr. P. Nagarajan for the help with animal husbandry. We deeply appreciate Prof. G. Ghosh, UCSD and Dr. R. Gokhale for critical comment on our work. Research in the PI’s laboratory was funded by DBT and NII-Core. MC thank DBT and AD thank DST-INSPIRE for research fellowships.

## Author Contribution

MC performed the animal experiments with help from BB; MC, TM, AD, and UAS, carried out the biochemical studies; MC performed the global gene analyses with guidance from SKB and SB, and BC carried out data analyses; SK, and AKS, provided human samples for MC, to examine under the guidance of VA and SB; MC and BC performed all other statistical tests; SB, MC, TM and BB designed the experiments; SB conceived the study, provided overall supervision and wrote the manuscript with MC.

## Reference

Allen, I.C., Wilson, J.E., Schneider, M., Lich, J.D., Roberts, R.A., Arthur, J.C., Woodford, R.-M.T., Davis, B.K., Uronis, J.M., Herfarth, H.H., et al. (2012). NLRP12 suppresses colon inflammation and tumorigenesis through the negative regulation of noncanonical NF-κB signaling. Immunity 36, 742–754.

Almaden, J. V., Tsui, R., Liu, Y.C., Birnbaum, H., Shokhirev, M.N., Ngo, K.A., Davis-Turak, J.C., Otero, D., Basak, S., Rickert, R.C., et al. (2014). A Pathway Switch Directs BAFF Signaling to Distinct NFκB Transcription Factors in Maturing and Proliferating B Cells. Cell Rep. 9, 2098–2111.

Andresen, L., Jørgensen, V.L., Perner, A., Hansen, A., Eugen-Olsen, J., and Rask-Madsen, J. (2005). Activation of nuclear factor kappaB in colonic mucosa from patients with collagenous and ulcerative colitis. Gut 54, 503–509.

Baeuerle, P.A., and Baltimore, D. (1988). IκB: A specific inhibitor of the NF-κB transcription factor. Science (80-.).

Banoth, B., Chatterjee, B., Vijayaragavan, B., Prasad, M.V.R., Roy, P., and Basak, S. (2015). Stimulus-selective crosstalk via the NF-κB signaling system reinforces innate immune response to alleviate gut infection. Elife 4, 1–56.

Basak, S., Kim, H., Kearns, J.D.J.D., Tergaonkar, V., O’Dea, E., Werner, S.L.S.L., Benedict, C.A.C.A., Ware, C.F.C.F., Ghosh, G., Verma, I.M.I.M., et al. (2007). A Fourth IκB Protein within the NF-κB Signaling Module. Cell 128, 369–381.

Basak, S., Shih, V.F.-S., and Hoffmann, A. (2008). Generation and Activation of Multiple Dimeric Transcription Factors within the NF- B Signaling System. Mol. Cell. Biol. 28, 3139–3150.

Boutaffala, L., Bertrand, M.J.M., Remouchamps, C., Seleznik, G., Reisinger, F., Janas, M., Bénézech, C., Fernandes, M.T., Marchetti, S., Mair, F., et al. (2015). NIK promotes tissue destruction independently of the alternative NF-κB pathway through TNFR1/RIP1-induced apoptosis. Cell Death Differ. 22, 2020–2033.

Brightbill, H.D., Suto, E., Blaquiere, N., Ramamoorthi, N., Sujatha-Bhaskar, S., Gogol, E.B., Castanedo, G.M., Jackson, B.T., Kwon, Y.C., Haller, S., et al. (2018). NF-κB inducing kinase is a therapeutic target for systemic lupus erythematosus. Nat. Commun. 9, 179.

Burkitt, M.D., Hanedi, A.F., Duckworth, C.A., Williams, J.M., Tang, J.M., O’Reilly, L.A., Putoczki, T.L., Gerondakis, S., Dimaline, R., Caamano, J.H., et al. (2015). NF-κB1, NF-κB2 and c-Rel differentially regulate susceptibility to colitis-associated adenoma development in C57BL/6 mice. J. Pathol. 236, 326–336.

Chatterjee, B., Banoth, B., Mukherjee, T., Taye, N., Vijayaragavan, B., Chattopadhyay, S., Gomes, J., and Basak, S. (2016). Late-phase synthesis of I B insulates the TLR4-activated canonical NF- B pathway from noncanonical NF- B signaling in macrophages. Sci. Signal. 9, ra120–ra120.

Choi, Y., Koh, S.-J., Lee, H.S., Kim, J.W., Gwan Kim, B., Lee, K.L., and Kim, J.S. (2015). Roxithromycin inhibits nuclear factor kappaB signaling and endoplasmic reticulum stress in intestinal epithelial cells and ameliorates experimental colitis in mice. Exp. Biol. Med. 240, 1664–1671.

Cohavy, O., Zhou, J., Ware, C.F., and Targan, S.R. (2005). LIGHT Is Constitutively Expressed on T and NK Cells in the Human Gut and Can Be Induced by CD2-Mediated Signaling. J. Immunol. 174, 646–653.

Dhar, A., Chawla, M., Chattopadhyay, S., Oswal, N., Umar, D., Gupta, S., Bal, V., Rath, S., George, A., Arimbasseri, G.A.A., et al. (2019). Role of NF-kappaB2-p100 in regulatory T cell homeostasis and activation. Sci. Rep. 9, 13867.

Eckmann, L., Nebelsiek, T., Fingerle, A.A., Dann, S.M., Mages, J., Lang, R., Robine, S., Kagnoff, M.F., Schmid, R.M., Karin, M., et al. (2008). Opposing functions of IKK during acute and chronic intestinal inflammation. Proc. Natl. Acad. Sci. 105, 15058–15063.

Franco, N.H., Correia-Neves, M., and Olsson, I.A.S. (2012). How “humane” is your endpoint?-refining the science-driven approach for termination of animal studies of chronic infection. PLoS Pathog.

Friedrich, M., Pohin, M., and Powrie, F. (2019). Cytokine Networks in the Pathophysiology of Inflammatory Bowel Disease. Immunity 50, 992–1006.

Grinberg-Bleyer, Y., Caron, R., Seeley, J.J., De Silva, N.S., Schindler, C.W., Hayden, M.S., Klein, U., and Ghosh, S. (2018). The Alternative NF-κB Pathway in Regulatory T Cell Homeostasis and Suppressive Function. J. Immunol. 200, 2362–2371.

Han, Y.M., Koh, J., Kim, J.W., Lee, C., Koh, S.-J., Kim, B., Lee, K.L., Im, J.P., and Kim, J.S. (2017). NF-kappa B activation correlates with disease phenotype in Crohn’s disease. PLoS One 12, e0182071.

Hoffmann, A., and Leung, T.H. (2003). Genetic analysis of NF- k B / Rel transcription factors de ® nes functional speci ® cities. EMBO 22, 5530–5539.

Jie, Z., Yang, J.-Y.Y., Gu, M., Wang, H., Xie, X., Li, Y., Liu, T., Zhu, L., Shi, J., Zhang, L., et al. (2018). NIK signaling axis regulates dendritic cell function in intestinal immunity and homeostasis. Nat. Immunol. 19, 1224–1235.

Jungbeck, M., Stopfer, P., Bataille, F., Nedospasov, S.A., Männel, D.N., and Hehlgans, T. (2008). Blocking lymphotoxin beta receptor signalling exacerbates acute DSS-induced intestinal inflammation—Opposite functions for surface lymphotoxin expressed by T and B lymphocytes. Mol. Immunol. 45, 34–41.

Karrasch, T., Kim, J.-S., Muhlbauer, M., Magness, S.T., and Jobin, C. (2007). Gnotobiotic IL-10 −/− ;NF-κB EGFP Mice Reveal the Critical Role of TLR/NF-κB Signaling in Commensal Bacteria-Induced Colitis. J. Immunol. 178, 6522–6532.

Kiesler, P., Fuss, I.J., and Strober, W. (2015). Experimental Models of Inflammatory Bowel Diseases. Cell. Mol. Gastroenterol. Hepatol. 1, 154–170.

Kotas, M.E., and Medzhitov, R. (2015). Homeostasis, Inflammation, and Disease Susceptibility. Cell.

Liu, J.Z., Van Sommeren, S., Huang, H., Ng, S.C., Alberts, R., Takahashi, A., Ripke, S., Lee, J.C., Jostins, L., Shah, T., et al. (2015). Association analyses identify 38 susceptibility loci for inflammatory bowel disease and highlight shared genetic risk across populations. Nat. Genet.

Liu, T., Zhang, L., Joo, D., and Sun, S.-C. (2017). NF-κB signaling in inflammation. Signal Transduct. Target. Ther. 2, 17023.

Love, M.I., Huber, W., and Anders, S. (2014). Moderated estimation of fold change and dispersion for RNA-seq data with DESeq2. Genome Biol.

Lyons, J., Ghazi, P.C., Starchenko, A., Tovaglieri, A., Baldwin, K.R., Poulin, E.J., Gierut, J.J., Genetti, C., Yajnik, V., Breault, D.T., et al. (2018). The colonic epithelium plays an active role in promoting colitis by shaping the tissue cytokine profile. PLOS Biol. 16, e2002417.

Macho-Fernandez, E., Koroleva, E.P., Spencer, C.M., Tighe, M., Torrado, E., Cooper, A.M., Fu, Y.-X., and Tumanov, A. V. (2015). Lymphotoxin beta receptor signaling limits mucosal damage through driving IL-23 production by epithelial cells. Mucosal Immunol. 8, 403–413.

Mikuda, N., Schmidt-Ullrich, R., Kärgel, E., Golusda, L., Wolf, J., Höpken, U.E., Scheidereit, C., Kühl, A.A., and Kolesnichenko, M. (2020). Deficiency in IκBα in the intestinal epithelium leads to spontaneous inflammation and mediates apoptosis in the gut. J. Pathol.

Mise-Omata, S., Kuroda, E., Niikura, J., Yamashita, U., Obata, Y., and Doi, T.S. (2007). A Proximal κB Site in the IL-23 p19 Promoter Is Responsible for RelA- and c-Rel-Dependent Transcription. J. Immunol. 179, 6596–6603.

Mitchell, S., Vargas, J., and Hoffmann, A. (2016). Signaling via the NFκB system. Wiley Interdiscip. Rev. Syst. Biol. Med. 8, 227–241.

Neurath, M.F., Pettersson, S., Meyer Zum Büschenfelde, K.-H., and Strober, W. (1996). Local administration of antisense phosphorothiate olignucleotides to the p65 subunit of NF–κB abrogates established experimental colitis in mice. Nat. Med. 2, 998–1004.

Ramakrishnan, S.K., Zhang, H., Ma, X., Jung, I., Schwartz, A.J., Triner, D., Devenport, S.N., Das, N.K., Xue, X., Zeng, M.Y., et al. (2019). Intestinal non-canonical NFκB signaling shapes the local and systemic immune response. Nat. Commun. 10, 660.

Savinova, O. V., Hoffmann, A., and Ghosh, G. (2009). The Nfkb1 and Nfkb2 Proteins p105 and p100 Function as the Core of High-Molecular-Weight Heterogeneous Complexes. Mol. Cell 34, 591–602.

Shibata, W., Maeda, S., Hikiba, Y., Yanai, A., Ohmae, T., Sakamoto, K., Nakagawa, H., Ogura, K., and Omata, M. (2007). Cutting Edge: The I B Kinase (IKK) Inhibitor, NEMO-Binding Domain Peptide, Blocks Inflammatory Injury in Murine Colitis. J. Immunol. 179, 2681–2685.

Shih, V.F.-S.S., Tsui, R., Caldwell, A., and Hoffmann, A. (2011). A single NFκB system for both canonical and non-canonical signaling. Cell Res. 21, 86–102.

Steinbrecher, K.A., Harmel-Laws, E., Sitcheran, R., and Baldwin, A.S. (2008). Loss of Epithelial RelA Results in Deregulated Intestinal Proliferative/Apoptotic Homeostasis and Susceptibility to Inflammation. J. Immunol. 180, 2588–2599.

Sun, S.-C. (2017). The non-canonical NF-κB pathway in immunity and inflammation. Nat. Rev. Immunol. 17, 545–558.

Tao, Z., Fusco, A., Huang, D.-B., Gupta, K., Young Kim, D., Ware, C.F., Van Duyne, G.D., and Ghosh, G. (2014). p100/IκBδ sequesters and inhibits NF-κB through kappaBsome formation. Proc. Natl. Acad. Sci. 111, 15946–15951.

Tsui, R., Kearns, J.D., Lynch, C., Vu, D., Ngo, K.A., Basak, S., Ghosh, G., and Hoffmann, A. (2015). IκBβ enhances the generation of the low-affinity NFκB/RelA homodimer. Nat. Commun. 6, 7068.

Upadhyay, V., and Fu, Y.-X. (2013). Lymphotoxin signalling in immune homeostasis and the control of microorganisms. Nat. Rev. Immunol. 13, 270–279.

Wullaert, A., Bonnet, M.C., and Pasparakis, M. (2011). NF-κB in the regulation of epithelial homeostasis and inflammation. Cell Res. 21, 146–158.

Yilmaz, Z.B., Kofahl, B., Beaudette, P., Baum, K., Ipenberg, I., Weih, F., Wolf, J., Dittmar, G., and Scheidereit, C. (2014). Quantitative Dissection and Modeling of the NF-κB p100-p105 Module Reveals Interdependent Precursor Proteolysis. Cell Rep. 9, 1756–1769.

Zaidi, D., and Wine, E. (2018). Regulation of Nuclear Factor Kappa-Light-Chain-Enhancer of Activated B Cells (NF-κβ) in Inflammatory Bowel Diseases. Front. Pediatr. 6, 1–9.

Zarnegar, B., Yamazaki, S., He, J.Q., and Cheng, G. (2008). Control of canonical NF-κB activation through the NIK-IKK complex pathway. Proc. Natl. Acad. Sci. U. S. A.

