## Supplementary file for "An epithelial *Nfkb2* pathway exacerbates intestinal inflammation by supplementing latent RelA dimers to the canonical NF-κB module"

### # Figure S1 – Figure S5;

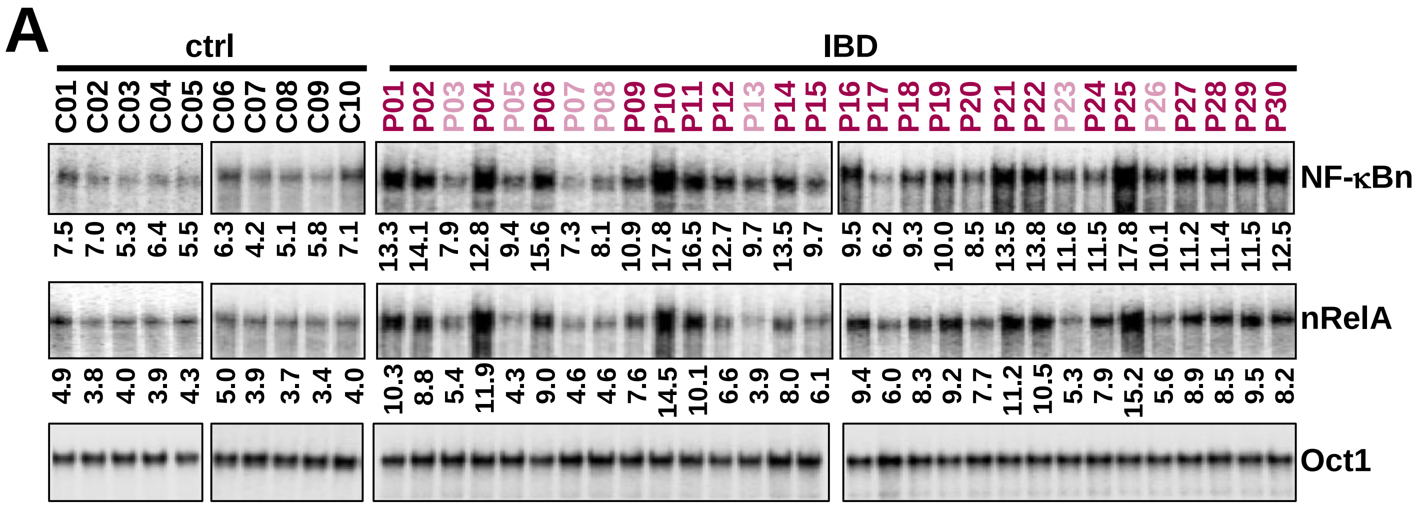


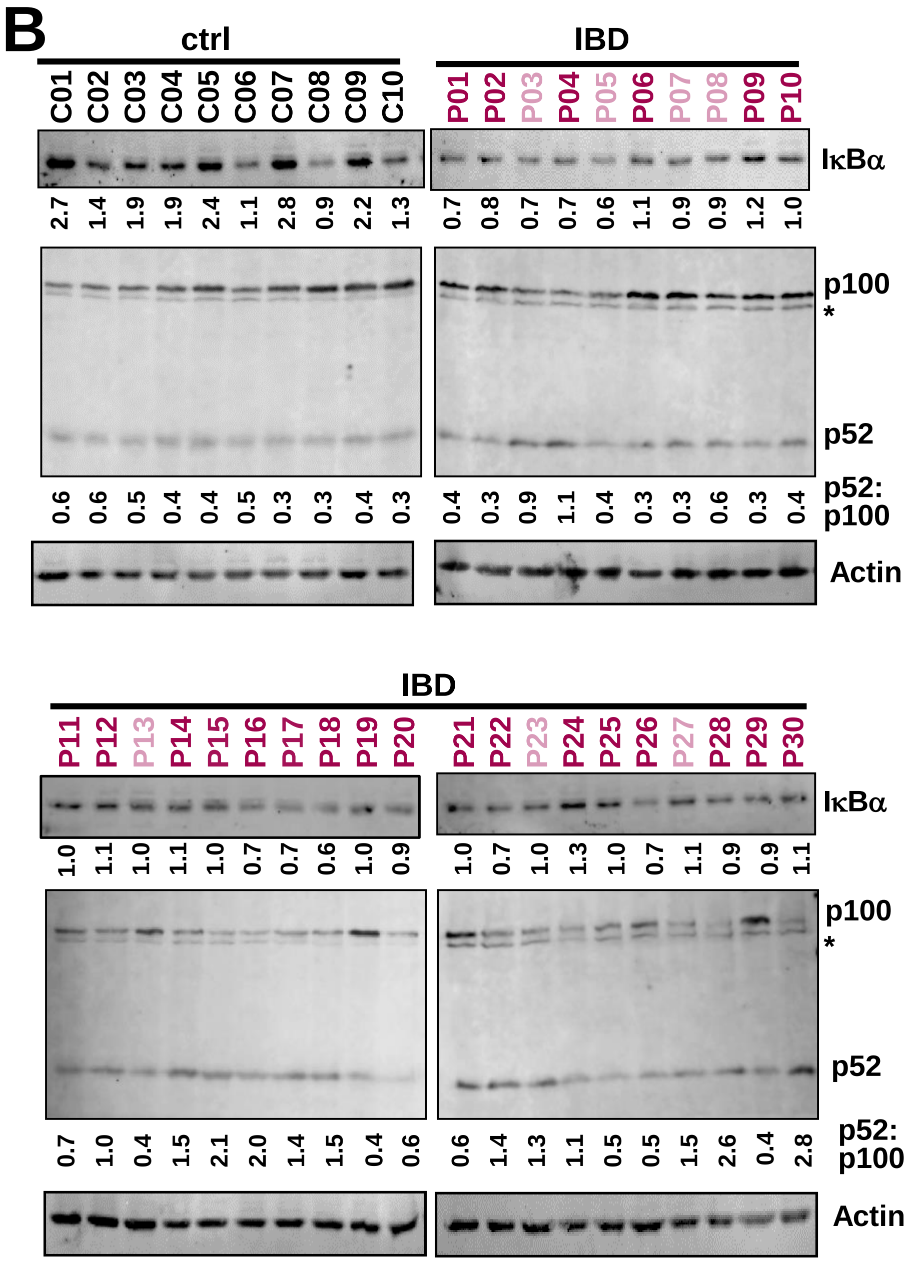


**Figure S1: Biochemical analyses of colonic tissues derived from human subjects. A.** EMSA revealing nuclear NF-κB DNA binding activity (NF-κBn, top panel) in colonic tissues from IBD patients. Controls signify non-IBD, hemorrhoids patients. Ablating RelB and cRel complexes using anti-RelB and anti-cRel antibodies, nuclear RelA DNA binding activities (nRelA) were examined in RelA-EMSA (middle panel). Oct1 DNA binding (bottom panel) served as a loading control. Signals corresponding to total NF-κBn and nRelA activities were quantified by densitometric analyses and presented below the respective lanes. Biopsies from a total of ten controls and thirty IBD patients were investigated. nRelA$_{high}$ and nRelA$_{low}$ IBD patients have been indicated with different font colors.
**B.** Immunoblot analyses revealing the abundance of IκBα (top panel), p52 as well

C

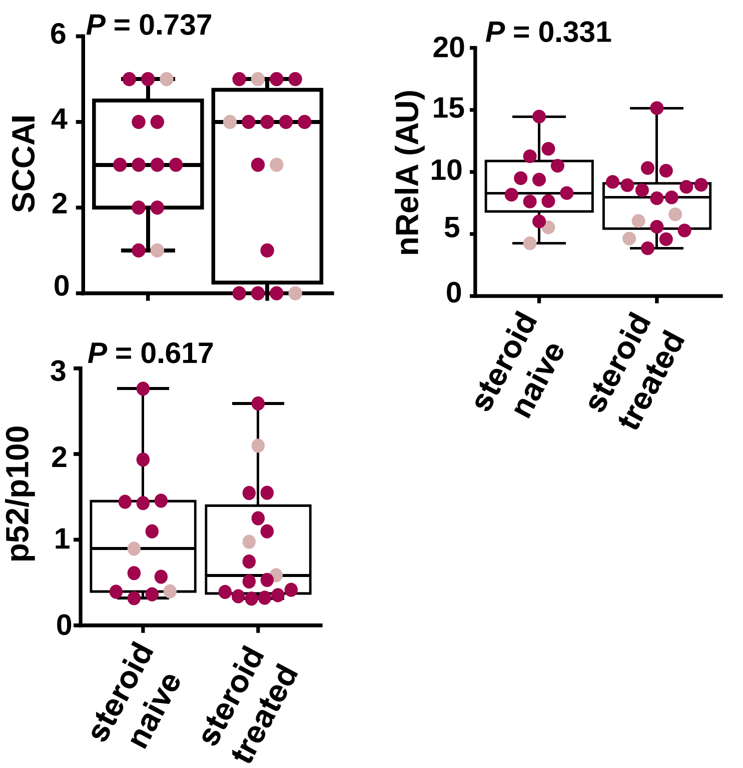


D

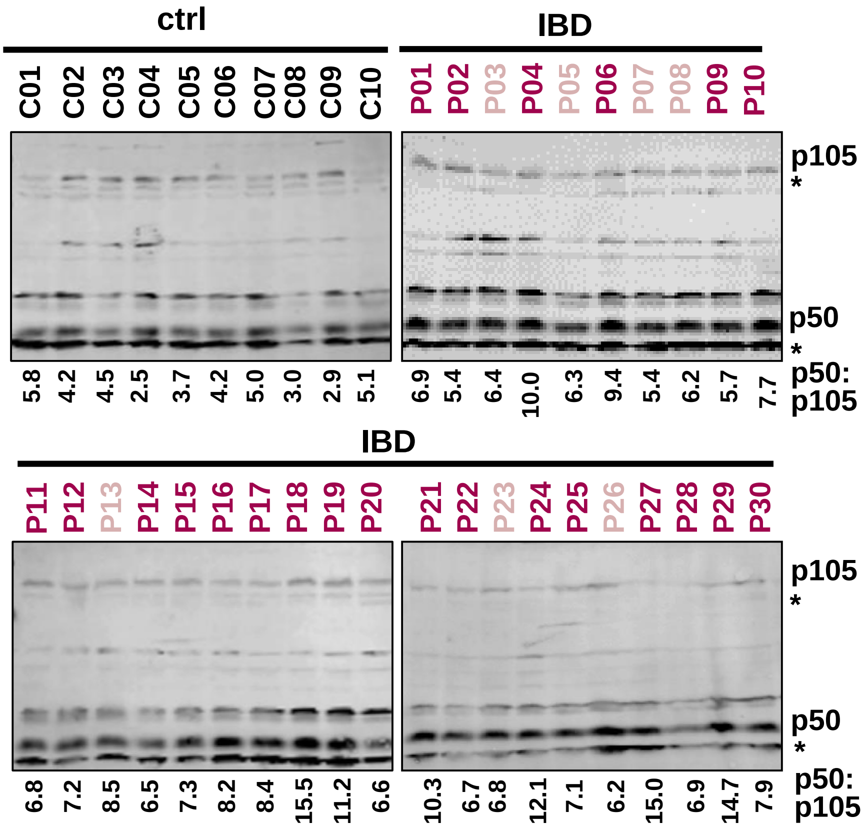


as p100 (middle panel) in whole-cell extracts obtained using colonic tissues from control individuals and IBD patients. Actin served as a loading control (bottom panel).Signals were quantified, the abundance of IκBα or the relative abundance of p52 to p100 (p52:p100) was determined and presented below the respective lanes. * indicates nonspecific protein bands ascertained using cell extracts obtained from WT or *Nfkb2*$^{-/-}$ mice.

**C.** Comparing steroid-naïve and steroid-treated patients for disease severity, nRelA activity and p100 processing. The cohort consisted of thirteen steroid-naïve and seventeen steroid-treated patients. Disease activity in UC was measured by Simple Clinical Colitis Activity Index (SCCAI). Disease activity status of one individual was unknown.

**D.** Immunoblot analyses was performed for detecting p50 and p105 in whole-cell extracts from controls and IBD patients. Signals were quantified and the relative abundance of p50 to p105 (p50:p105) was determined and presented below the respective lanes. * indicates nonspecific protein bands ascertained using cell extracts obtained from WT or *Nfkb1*$^{-/-}$ mice.

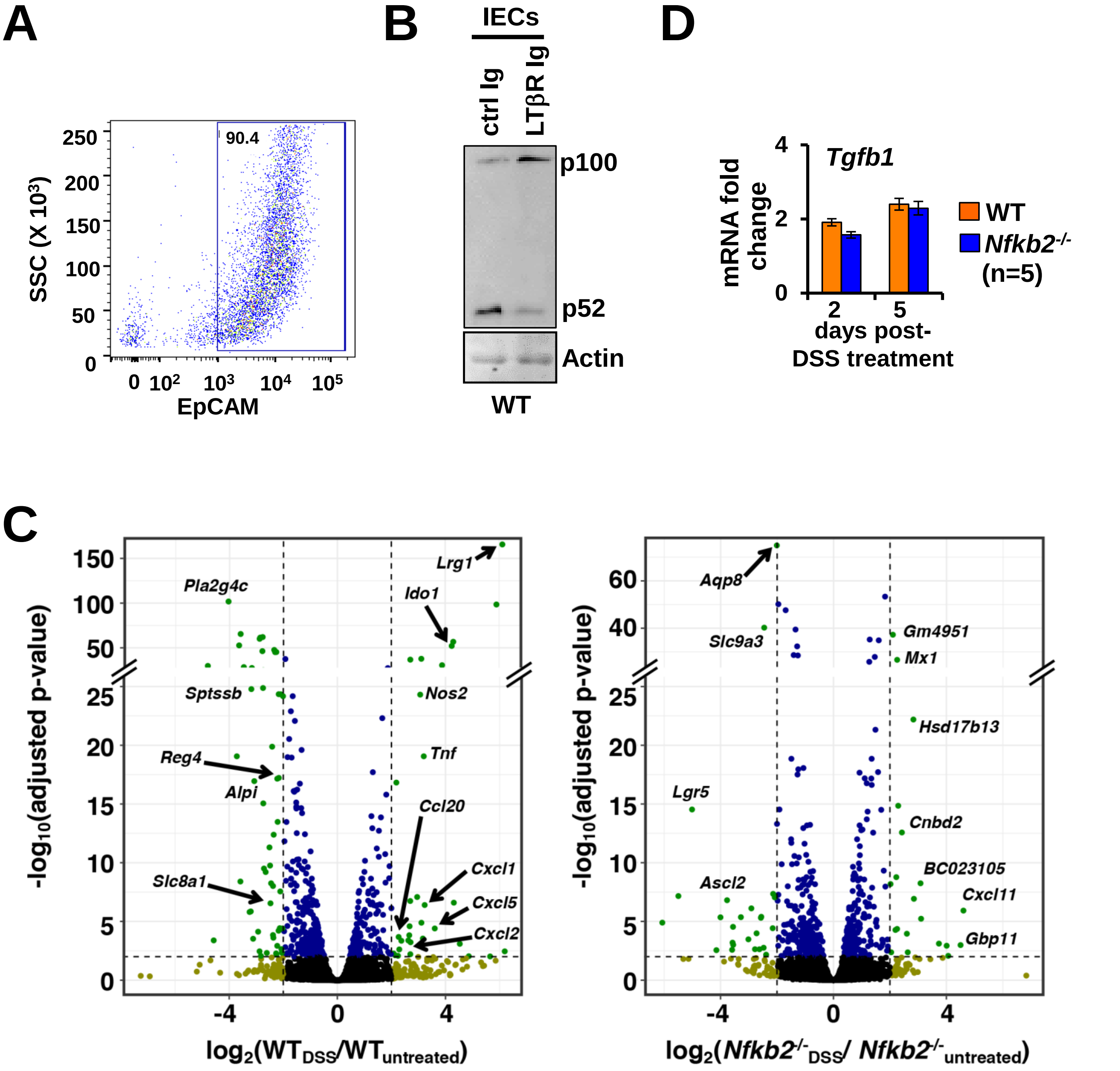


**Figure S2: Investigating the *Nfkb2* pathway in IECs derived from colitogenic mice. A.** FACS plots revealing enrichment of EpCAM+ cells in our mouse IEC preparation. **B.** Immunoblot of whole-cell extracts derived from IECs from WT mice administered with either control-Ig or LTβR-Ig 24h prior to tissue collection. **C.** Volcano plots depicting the fold change in the mRNA level in IECs upon 48h of DSS treatment of WT (left) or *Nfkb2*$^{-/-}$ (right) mice. The global gene expression data represents three biological replicates. The horizontal dashed line represents p-value = 0.01. The green dots represent genes, whose expressions were at least four fold different in a stastistically significant manner. **D.** Gene expression analyses revealing the abundance of indicated mRNAs in IECs derived from WT and *Nfkb2*$^{-/-}$ mice administered with 2.5% DSS for two or five days. mRNA fold change values were calculated in relation to corresponding untreated mice (n=5). Data represent means ± SEM. The statistical significance was determined using two-tailed Student's t-test. ** *P*<0.01.

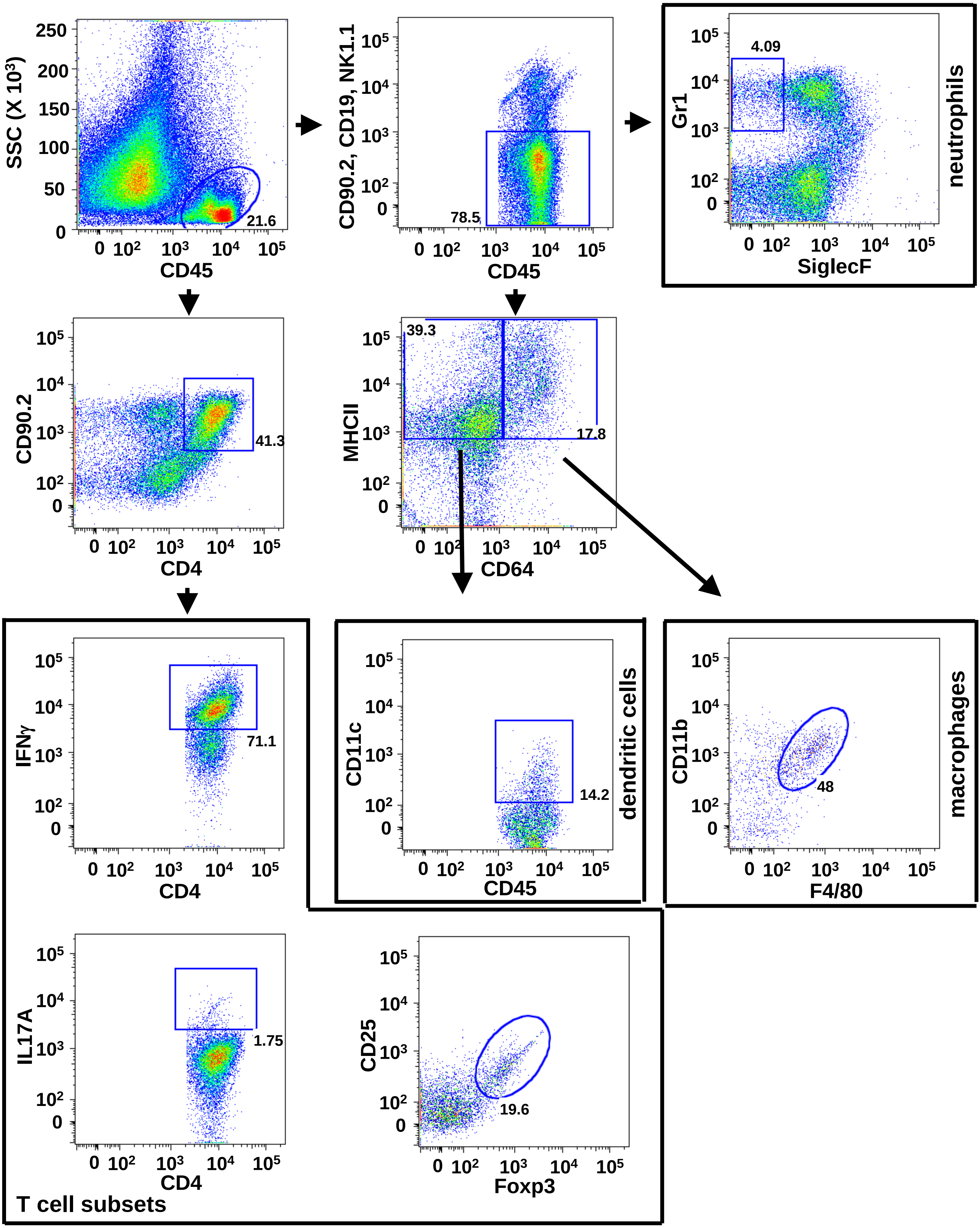


**Figure S3: Investigating immune cells in the colitogenic gut of WT and *Nfkb2*$^{-/-}$ mice.** Representative FACS plots showing the general gating strategy for analysing the frequencies of neutrophils, macrophages, dendritic cells, and variout T cell subsets in the lamina propria. The data represent WT mice treated with DSS for five days. Similarly, frequencies of these cells were also measured in DSS-treated *Nfkb2*$^{-/-}$ mice.

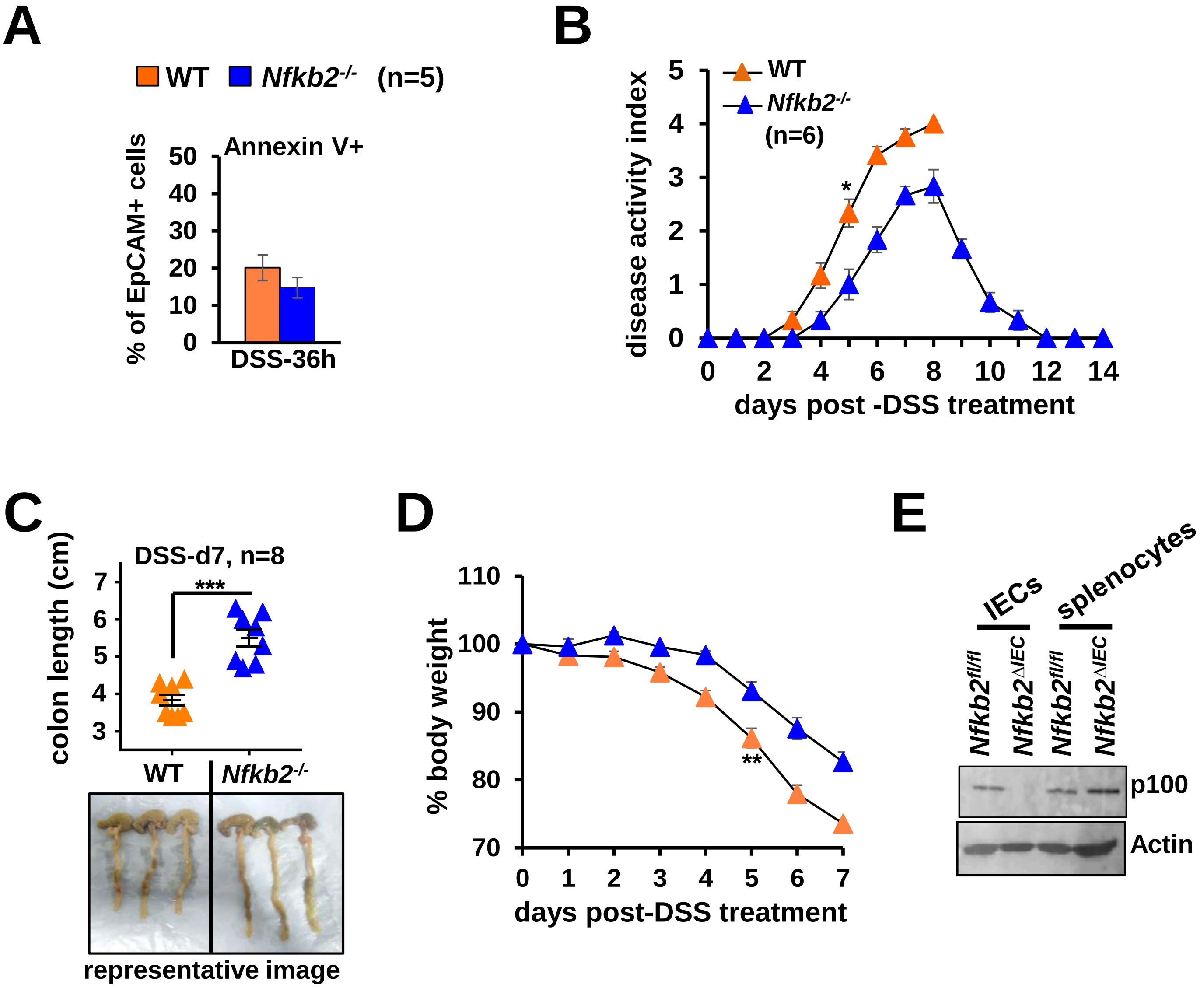


**Figure S4: Comparing WT and *Nfkb2*$^{-/-}$ mice for colitogenic phenotypes. A.** WT and *Nfkb2*$^{-/-}$ mice were administered with 2.5% DSS for 36h, subequently IECs were collected and examined for the presence of Annexin V+ apoptotic cells; the data has been corrected for corresponding basal apoptosis observed in untrated mice. Alternatly, mice were treated with DSS for seven days and evaluated for the disease activity **(B)**, colon length **(C)** and and bodyweight changes **(D)**. In the case of WT, disease activity measurements were discontinued after nine days because of morbidity and mortality **(B)**. Colon length was measured at day seven from the onset of DSS treatment; representative images of colons derived from DSS-treated WT or *Nfkb2*$^{-/-}$ mice have also been shown **(C)**. Bodyweight changes were scored in a time course **(D)**. **E.** Representative immunoblot revealing the abundance of p100/*Nfkb2* in IECs and spleenocytes derived from *Nfkb2*$^{fl/fl}$ and *Nfkb2*$^{\Delta IEC}$ mice. Statistical significance was determined by two-tailed Student's t-test. ***$P$ < 0.001; **$P$ < 0.01; *$P$ < 0.05.

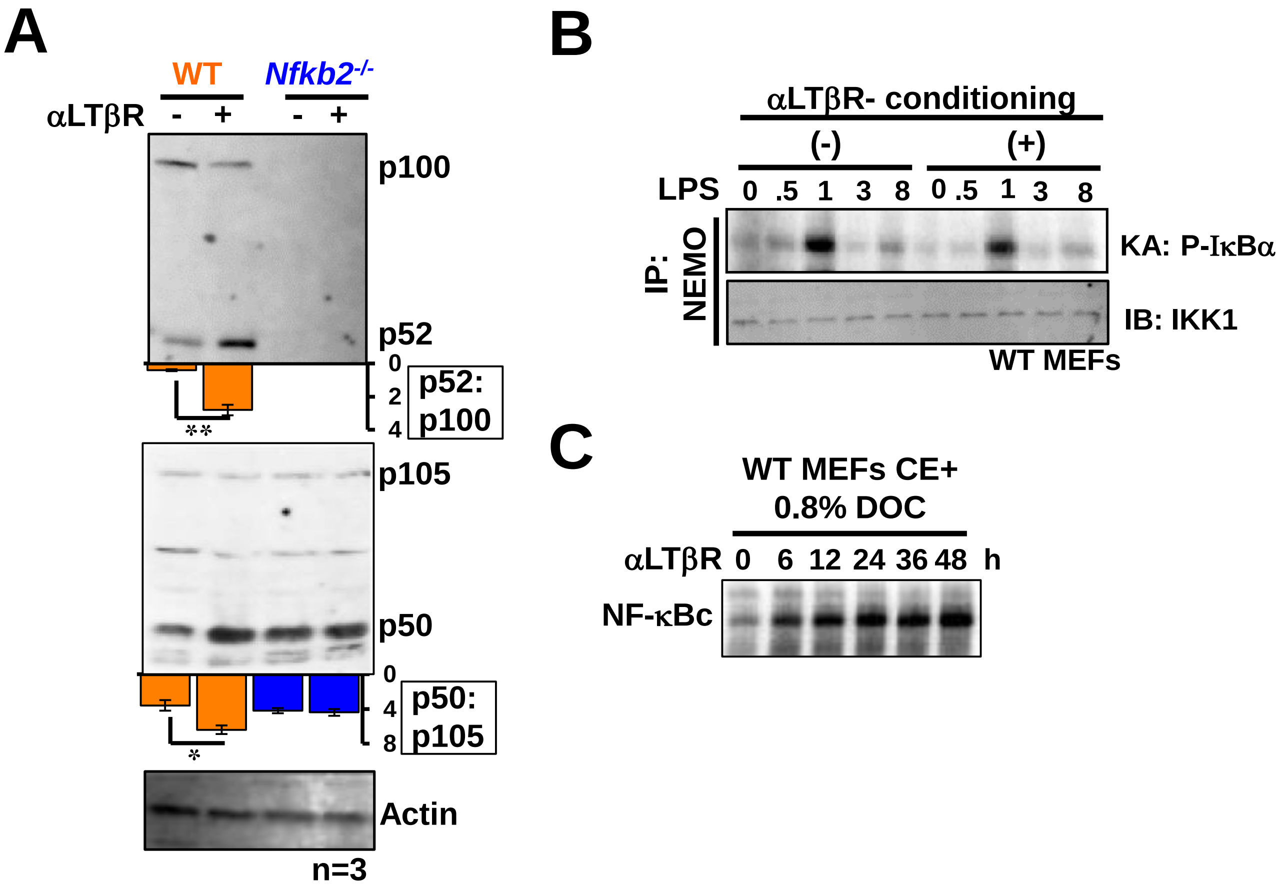


**Figure S5: Investigating crosstalks between LTβR-*Nfkb2* signaling and the canonical NF-κB pathway.** **A.** Immunoblot revealing processing of p100 and p105 into p52 and p50, respectively, in LTβR stimulated WT or *Nfkb2*$^{-/-}$ MEFs. Cells were stimulated for 36h using 0.1µg/ml of αLTβR. Data represent three experimental replicates. Signals were quantified, the relative abundance of p52 to p100 (p52:p100) or p50 to p105 (p50:p105) was determined and presented below the respective lanes. **B.** Kinase assay revealing the activation of the NEMO-IKK2 complex in MEFs upon LPS treatment. Cells were either left untreated or treated with 0.1µg/ml of αLTβR agonistic antibody, which activates the noncanonical pathway, for 36h before being subjected to LPS stimulation. Subsequently, NEMO co-immunoprecpipitates obtained from these cells were incubated with recombinant GST-IκBα for scoring the NEMO-IKK2 activity. IKK1 co-immunoprecipitated with NEMO was probed as loading control. **C.** EMSA revealing gradual accumulation of latent NF-κB complexes in the cytoplasm of cells subjected to αLTβR stimulation (0.1µg/ml of αLTβR). Cytoplasmic extracts were treated with deoxycholate before being subjected to EMSA for unmasking latent NF-κB DNA binding activities.
